# The context-dependent, combinatorial logic of BMP signaling

**DOI:** 10.1101/2020.12.08.416503

**Authors:** Heidi Klumpe, Matthew A. Langley, James M. Linton, Christina J. Su, Yaron E. Antebi, Michael B. Elowitz

**Affiliations:** Division of Biology and Bioengineering, California Institute of Technology, Pasadena, CA 91125, USA; Division of Chemistry and Chemical Engineering, California Institute of Technology, Pasadena, CA 91125, USA; Department of Molecular Genetics, Weizmann Institute of Science, Rehovot 76100, Israel; Howard Hughes Medical Institute, Chevy Chase, MD 20815, USA; Department of Applied Physics, California Institute of Technology, Pasadena, CA 91125, USA

**Keywords:** bone morphogenetic protein, BMP, signaling pathways, cell context, combinatorial signaling, promiscuous receptor-ligand interactions, pairwise interaction analysis

## Abstract

Cell-cell communication systems typically comprise families of ligand and receptor variants that function together in combinations. Pathway activation depends in a complex way on which ligands are present and what receptors are expressed by the signal-receiving cell. To understand the combinatorial logic of such a system, we systematically measured pairwise Bone Morphogenetic Protein (BMP) ligand interactions in cells with varying receptor expression. Ligands could be classified into equivalence groups based on their profile of positive and negative synergies with other ligands. These groups varied with receptor expression, explaining how ligands can functionally replace each other in one context but not another. Context-dependent combinatorial interactions could be explained by a biochemical model based on competitive formation of alternative signaling complexes with distinct activities. Together, these results provide insights into the roles of BMP combinations in developmental and therapeutic contexts and establish a framework for analyzing other combinatorial, context-dependent signaling systems.

## Introduction

Cell communication pathways such as BMP, Wnt, and FGF play pivotal roles in normal development, disease, therapeutics, and regenerative medicine (Salazar, Gamer, and Rosen 2016; Nusse 2005; Ornitz and Itoh 2015). A striking feature of these pathways is their use of families of homologous ligand variants. Within each pathway, multiple ligands typically function together in combinations (Antebi et al. 2017; Alok et al. 2017; Kadam, Ghosh, and Stathopoulos 2012), and the effect of any given ligand can vary dramatically with cell or tissue context (Chen et al. 2013; Kadam et al. 2009; Fortin et al. 2005; Eubelen et al. 2018; Verkaar and Zaman 2010). Despite the prevalence of these features, and extensive information about the biochemical interactions and developmental roles of particular ligands, we lack a unified description of how ligands signal in combinations and across biological contexts. A clearer view of this structure could allow more predictive control of these pathways in natural and synthetic contexts.

A prime example of such combinatorial and contextual ligand activity can be seen in the Bone Morphogenetic Protein (BMP) pathway. This pathway comprises ten major ligand variants, as well as four Type I and three Type II receptors subunits (Figure 1A). Distinct combinations of these components regulate the development of diverse tissues including skeleton, kidney, eye, and brain (Salazar, Gamer, and Rosen 2016; Godin, Robertson, and Dudley 1999; Huang et al. 2015; Butler and Dodd 2003). Interestingly, a pair of ligands can exhibit either redundant or distinct roles depending on context. For example, BMP9 can replace the essential role of BMP10 in proper vasculature formation, but not in the developing heart (Chen et al. 2013). Similarly, BMP4 can replace BMP7 in the developing kidney, but not in the developing eye (Oxburgh et al. 2005). In addition to variation across tissue context, BMP ligands also have non-overlapping roles when signaling in combinations, such as observed in the developing joint (Bandyopadhyay et al. 2006), and can combine with each other non-additively (Piscione et al. 1997; Ying, Qi, and Zhao 2001; Klammert et al. 2015; Antebi et al. 2017).

**Figure 1:**
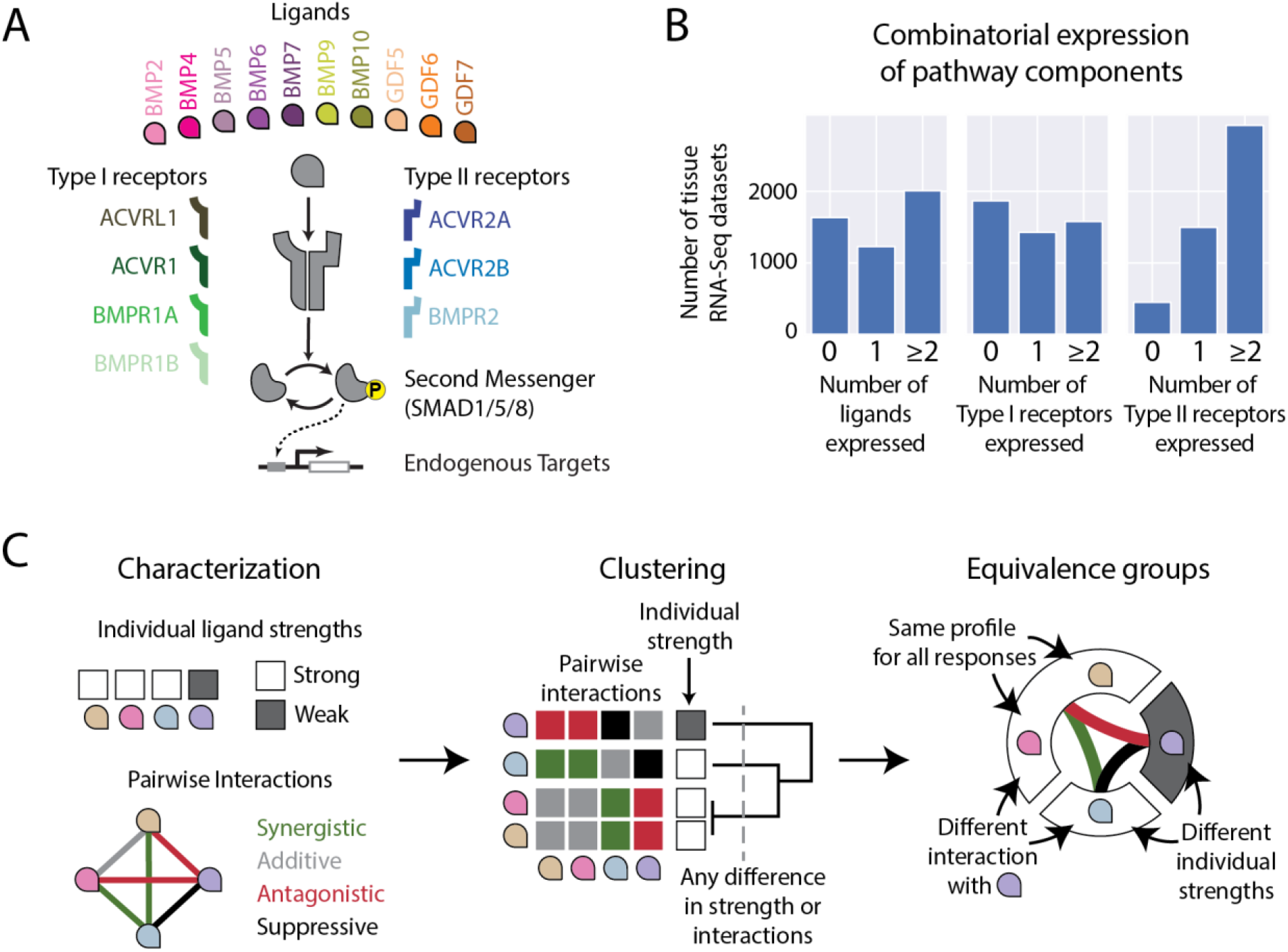
BMP ligands can be classified by their pairwise interactions. (A) BMP ligands (colored petal shapes) bind Type I and Type II receptors to form signaling complexes. Active signaling complexes phosphorylate SMAD1/5/8 transcription factors, which translocate to the nucleus to activate endogenous gene targets. Promiscuous ligand-receptor interactions enable the formation of a wide variety of potential signaling complexes. The dimeric nature of the ligands allows each to recruit up to two Type I and two Type II receptors simultaneously. (B) Bulk RNA-Seq datasets of various mouse tissues, collected in (Lachmann et al. 2018), show that BMP ligands, Type I receptors, and Type II receptors are often expressed (i.e. FPKM > 0) as combinations (i.e. 2 or more). (C) Systematic measurements of individual and pairwise responses (left) can cluster BMP ligands into equivalence groups (middle). Ligands in the same equivalence group interact in similar ways (synergistic, additive, antagonistic, or suppressive) with other ligands and have the same individual strength (right).

Thus far, the primary framework for describing BMP ligand differences is variation in ligand-receptor affinity. Ligand-receptor affinities govern the formation of signaling complexes, each of which contains a dimeric ligand bound to two Type I and two Type II receptor subunits Figure 1A). These signaling complexes phosphorylate SMAD1/5/8 effector proteins (as well as activate other non-canonical targets) to regulate target genes. Ligands are often grouped by their preferred Type I and Type II receptors to summarize their functional differences (Shimasaki et al. 2004; C. J. David and Massagué 2018). However, receptor preferences cannot explain key functional differences between ligands. For instance BMP9 and BMP10 exhibit similar receptor preferences (L. David et al. 2007) but behave non-equivalently, as mentioned above (Chen et al. 2013). Moreover, BMP receptors are expressed in combinations (Figure 1B), compete with one another for binding to ligands, and generate complex functional responses to ligand combinations, all of which make it difficult to predict the distribution of signaling complexes, and thereby the overall pathway activity, from qualitative affinity preferences alone (Antebi et al. 2017).

Therefore, despite extensive molecular characterization, we cannot in general say how specific ligands will function in combinations, why these combinatorial effects vary with context, and what molecular features are necessary to generate the combinatorial and contextual behavior of BMP ligands. Addressing these questions would advance our understanding of the role of BMP signaling components in a variety of skeletal, respiratory, and brain diseases (Salazar, Gamer, and Rosen 2016; Valer et al. 2019; Cunha and Pietras 2011) and aid the selection of therapeutic BMPs to selectively induce bone growth in orthopedics and oral surgery (Cicciù et al. 2014; Carragee, Hurwitz, and Weiner 2011; Seeherman et al. 2019). More generally, it could enable more precise control of cell fate decisions for therapeutic applications and provide insight into the overall logic of cell-cell communication systems.

To obtain a systematic understanding of context-dependent ligand integration, we turned to pairwise interaction analysis, an approach that has provided powerful insights into the structure of other combinatorial biological systems. In this approach, one classifies pairs of perturbations as additive (neutral), positive (synergistic), or negative (antagonistic) depending on whether the perturbations are stronger or weaker in combination than expected from their individual effects. Pairwise analysis of mutations has revealed the structure of gene modules and protein functions (Schwikowski, Uetz, and Fields 2000; Segrè et al. 2005; Costanzo et al. 2016). Similarly, pairwise analysis of drug interactions successfully classified antibiotics into groups with similar biochemical mechanisms of action (Yeh, Tschumi, and Kishony 2006). We reasoned that pairwise analysis of BMP ligand combinations could be useful in two ways: First, it could provide a systematic map of how ligands combine to determine pathway activation (Figure 1C). Second, it could reveal how that combinatorial structure itself changes between different cell contexts. In this way, pairwise analysis of ligand combinations could provide a general way to understand systems that are both combinatorial and contextual.

With that in mind, we analyzed pairwise combinations of ten of the most important BMP ligands across multiple receptor contexts. (Analogous to the convention for drug interactions, we will use the term “ligand interactions” to refer to non-additive responses to ligand combinations, without implying direct molecular interaction between the ligands.) We first show how this pairwise analysis classifies BMP ligands into equivalence groups, defined such that ligands in the same group exhibit a similar strength and profile of effective interactions with other ligands (Figure 1C). This analysis reveals that the ten ligands can be organized into a smaller number of equivalence groups, and introduces a compact, useful representation for complex multi-ligand systems. (Ligand equivalence groups, defined here, are distinct from equivalence groups in development, which describe groups of cells with the same fate potential (Greenwald and Rubin 1992).) Next, we turn to the question of how ligand interactions vary with cell context, measuring ligand interactions using cells with differing receptor expression profiles. We find that receptor expression plays a powerful, non-intuitive role in shaping combinatorial ligand responses, and that receptor context effects can explain previous observations in development. Finally, to understand how the observed combinatorial ligand responses could emerge from underlying molecular features of the pathway, we fit the data to a mathematical model of competitive ligand-receptor interactions. We find that contextual multi-ligand BMP responses can emerge from the interplay between the affinities that govern signaling complex formation and the specific phosphorylation activities of the resulting complexes. Together, these results reveal the combinatorial and contextual logic of BMP signaling, and provide a framework for understanding other pathways that similarly integrate multi-ligand inputs with promiscuous protein-protein interactions.

## Results

### Quantitative dose-response measurements reveal ligand-specific features

Analyzing ligand combinations first requires an understanding of how the BMP pathway responds to ligands individually. To that end, we identified a core set of BMP ligands for our study from the larger set of TGF-β superfamily ligands (Weiss and Attisano 2013), focusing on ten homodimeric BMP ligands that play pivotal roles in development and are known to activate SMAD1/5/8 (Figure 1A) (Hinck, Mueller, and Springer 2016). Next, we constructed a BMP reporter cell line (Figure S1A). As a base cell line, we selected NAMRU mouse mammary gland (NMuMG) epithelial cells, which can respond to BMP signals without differentiating (Piek et al. 1999) and expresses five of the seven BMP receptors, including ACVR1, ACVR2A, ACVR2B, BMPR1A, and BMPR2 (Antebi et al. 2017). We integrated a construct encoding a fusion Histone 2B (H2B)-mCitrine protein controlled by a synthetic SMAD1/5/8-activated BMP responsive element (BRE) (Korchynskyi and ten Dijke 2002). The resulting stable cell line exhibited unimodal, dose-dependent distributions of YFP fluorescence levels in response to varying concentrations of BMP ligands (Figure S1B), allowing quantitative readout of pathway activity and ligand dose responses (Figure S1C).

The ligands exhibited a broad range of dose-response characteristics (Figure S1D). Seven ligands strongly activated the pathway but did so with varying saturating levels, concentrations for half-maximal activation (EC_50_), and logarithmic sensitivities, quantified by a fitted Hill coefficient. The EC_50_ ranged over three orders of magnitude among these ligands (Figure S1E) while the Hill coefficient varied between values of one and two (Figure S1F). By contrast, the Relative Ligand Strength (RLS), or the pathway activity at saturating concentrations normalized to the activity of the strongest ligand, could be quantified for all 10 ligands and fell into three distinct tiers (Figure S1G). In the lowest tier, GDF5, GDF6, and GDF7 barely activated the reporter above background. However, these ligands did activate the pathway following ectopic expression of BMPR1B (Figure S6E), a receptor not expressed by NMuMG cells and required for GDF signaling (Nishitoh et al. 1996).

### Ligands exhibit diverse combinatorial profiles

While the single-ligand dose responses identified ligands of equivalent strength (quantified here as RLS), they were not sufficient to classify ligand equivalence in combinations. To systematically probe combinatorial responses, we measured the responses to all 45 pairs of the 10 ligands (Figure 2A), as well as each ligand alone or paired with itself. The interactions of each ligand pair were observed at nine distinct ratios, using concentrations spanning the ligand’s dynamic range (Figure 2A, inset). A robotic liquid-handling system enabled measurements of multiple ligand combinations in a single experiment (STAR Methods), and responses were quantified by flow cytometry after 24h incubation with the ligands (Figures 2B-E).

**Figure 2:**
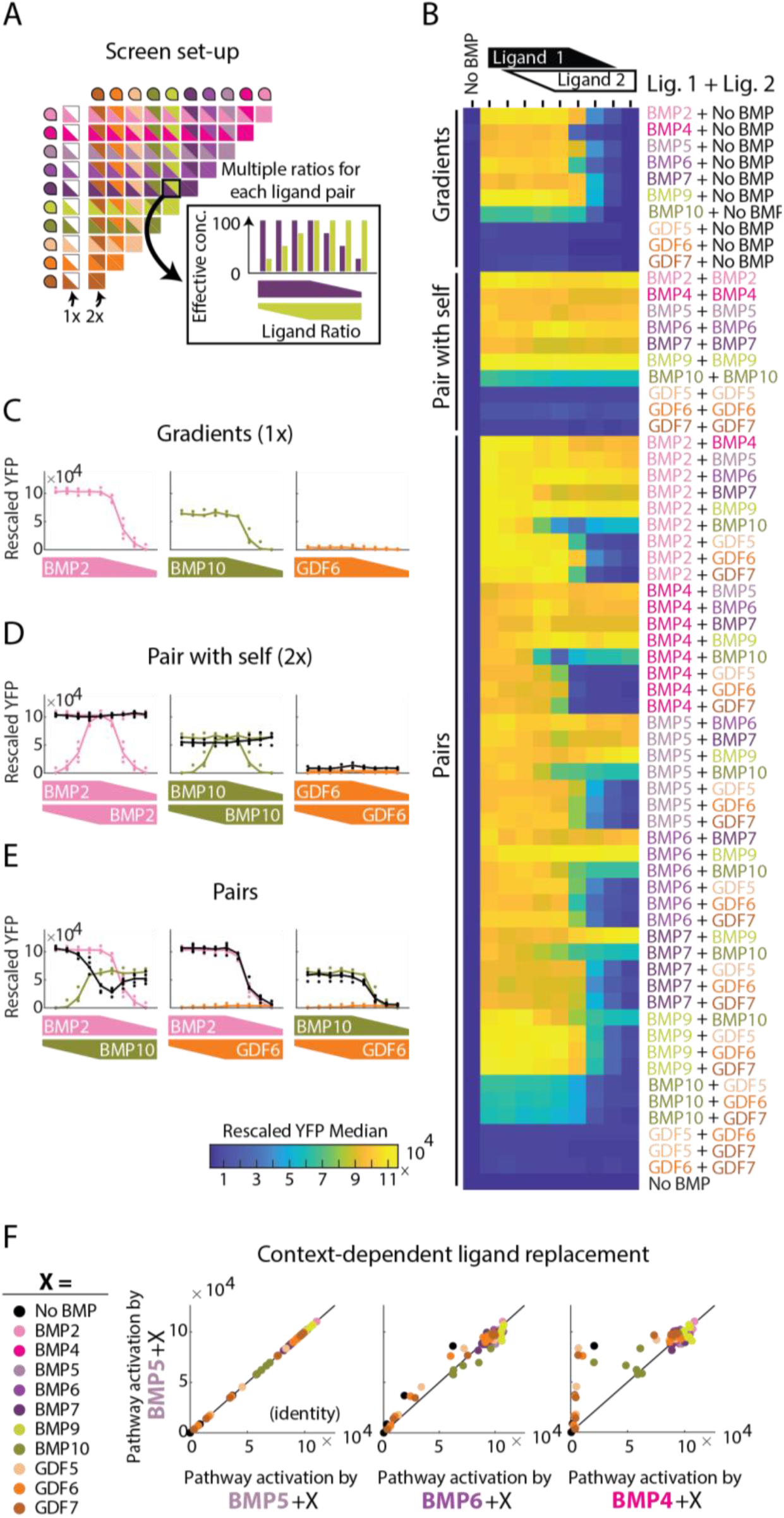
Pairwise titration of relative ligand concentrations reveals ligand interactions. (A) Fully classifying ligand equivalence groups requires analysis of all possible pairs, as well as measurements of individual activity (1X) and each ligand paired with itself (2X) as a positive control. Responses in each category are sampled at multiple ratios (inset), with at least one ligand very near its saturating concentration in each ratio. (B) The full dataset includes measurements for all possible gradients, pairs, and pairs with self, which are shown as each row of the heatmap. The columns correspond to concentration ratios, with the leftmost column reserved for a no BMP control. Medians of four biological repeats are shown with heatmap colors. (C) Sample gradients of BMP2, BMP10, and GDF6 show diversity of individual strength. Dots are four biological replicates, and the line connects the median value for each concentration. The ligand concentrations are high and saturating on the left, and decrease from left to right, as shown by the trapezoid height on the x-axis. (D) Ligands paired with themselves show purely additive relationships between ligands of the same strength. Colored dots and lines summarize biological replicates and median for the ligand of the indicated color, whereas the black line shows the pair indicated on the x-axis, either BMP2, BMP10, or GDF6 paired with itself for multiple concentration ratios. (E) All possible pairs of three ligands (BMP2, BMP10, and GDF6) show three types of pairwise interactions. Responses to pairs can be lower than both individual ligands (left), exactly equal to stronger of the two (middle), or interpolate between the two (right). As in C, the colored dots and lines show individual ligand responses, whereas black is the response to the pair. The concentration ratios are shown using the same trapezoids on the x-axis. (F) Responses to BMP5 paired with another ligand (“X”, colors) are plotted against the response to other ligands (BMP5, BMP6, or BMP7) also paired with X. Since all pairs are sampled at seven ratios, there are seven dots for each X (i.e. color). The first two plots show that BMP6 is a nearly equivalent replacement for BMP5, as BMP5+X closely resembles BMP6+X (i.e. dots are close to the y=x line). By contrast, BMP4 produces much lower activation than an equivalent amount of BMP5, in the presence of BMP10, GDF5, GDF6, or GDF7. Pathway activation is the median of four biological repeats, quantified by flow cytometry.

Comparing two ligand’s activities across combinatorial conditions summarizes their ability to produce equivalent pathway output (quantified here with YFP fluorescence) in the presence of another ligand. For example, consider BMP4, BMP5,and BMP6, which are all strong activators with RLS close to 1. By definition, replacing BMP5 with itself produces equivalent responses across all conditions (Figure 2F, left). Replacing BMP5 with BMP6 also produced very similar pathway activity, in the context of any other ligand or ratio, indicating that these two ligands behave equivalently in this context (Figure 2F, middle). By contrast, replacing BMP5 with an equivalent dose of BMP4 changed signaling activity in the presence of BMP10 or any of the GDFs (Figure 2F, right). Thus, ligands that exhibit similar RLS (strength) are not necessarily equivalent, due to the differences in their pairwise interactions with other ligands.

### Ligand interactions reveal BMP equivalence groups in NMuMG cells

Because differences in pairwise interactions are key to determining ligand equivalence, we sought a metric to quantitatively classify the type and strength of ligand interactions. Common drug interaction metrics, such as Loewe or Bliss, and epistasis metrics do not precisely map onto ligand interactions (Loewe 1928; Bliss 1939; Russ and Kishony 2018; Yeh, Tschumi, and Kishony 2006), so we defined the Interaction Coefficient (IC) (Figures 3A, S2A, STAR Methods). The IC is designed to map qualitatively different interaction types and magnitudes onto a single linear scale by comparing the combined and individual effects of two ligands. This scale separates qualitatively different behaviors into different IC regimes and includes both linear (IC=1) and saturated (IC=0) additivity, as well as negative interactions (IC<0) and positive synergy (IC>1). The resulting scale can be written as a set of piecewise-linear equations (Figure S2A), which correctly classify distinct behaviors (Figure S2B).

**Figure 3:**
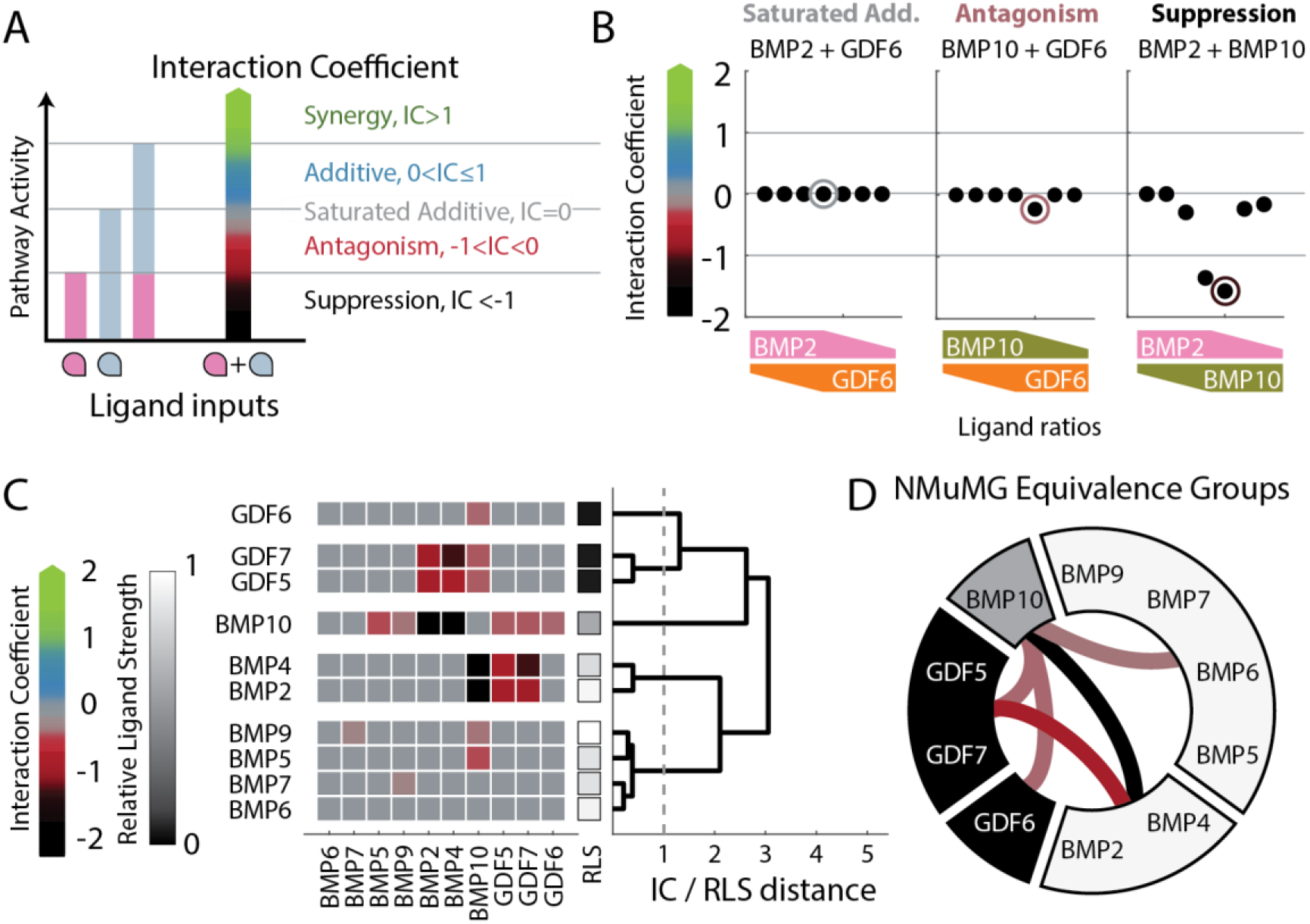
Antagonism and suppression define BMP equivalence groups in NMuMG cells. (A) The Interaction Coefficient (IC) classifies different regimes of pairwise responses by comparison to three references (horizontal lines): the two individual ligand responses, *f(A)* and *f(B)*, and the sum of those responses, *f(A) + f(B)*. The behavior type and strength for a given pairwise response, *f(A+B)*, is indicated by its IC value and associated color map: black for suppressive, red for antagonistic, gray for saturated additive, blue for additive, and green for synergistic. IC values linearly interpolate between reference points. Though IC has no upper bound, values rarely exceed 2, which is the upper limit shown on all plots. See Figure S1B and STAR Methods for formula. (B) IC values for pairs of BMP2, BMP10, and GDF6 (as in Figure 2E) vary across pairs and multiple concentration ratios. Each pair exhibits a distinct response type. Circled IC values indicate which ratio was used to classify the pair, and the circle’s color indicates the interaction type. Trapezoids on x-axis indicate concentration ratios, analogous to inset in A. (C) Hierarchical clustering of IC and RLS values in NMuMG cells, computed from the medians of four biological replicates, reveals five distinct equivalence groups, defined at a distance of 1 to separate ligands with similar pairwise and individual strengths (STAR Methods). The square colors indicate either the ligand’s IC value (see colormap in B) when paired with another ligand or the RLS grayscale for its individual strength, as indicated by x-tick labels. The dendrogram on the right-hand side shows complete-linkage clustering of Euclidean distance between ligand features. The RLS and IC values are weighted such that a distance of 1 indicates a significant difference in either individual strength or a difference of one pairwise interaction between two ligands (STAR Methods). (D) BMP ligand equivalence in NMuMG cells, as classified by hierarchical clustering in E, comprises five equivalence groups. Wedge grayscale value shows approximate RLS value of the contained ligands, while connecting line colors represent pairwise interactions between ligand groups. Groups without a linking edge interact additively. Ligands with similar individual strengths can be grouped differently based on their pairwise interactions. See also Figures S1, S2.

Despite its different definition, IC’s basic classifications mirror those of drug interactions, such as suppressive interactions (Yeh, Tschumi, and Kishony 2006; Bollenbach et al. 2009). These behaviors are also related to previously observed two-ligand response types, with balance, additive, ratiometric, and imbalance functions producing synergistic, additive, antagonistic, and suppressive interaction coefficients, respectively (Antebi et al. 2017). Moreover, like other interaction metrics, the IC value depends on the specific ligand concentrations for which it is computed, and is close to 1 at small enough concentrations of both ligands. Therefore, to characterize a ligand pair, we select the IC value of largest absolute value over all sampled concentration ratios (Figure 3B), as pairwise interactions are most evident at certain ligand ratios and with total concentrations close to saturation (Figure S2B). However, due to possible undersampling of concentration ratios and the unknown dynamic ranges of weakly activating ligands, these values may be underestimates of the strength of non-additive interactions.

Nonetheless, the interaction metric (IC) revealed an array of interaction types within NMuMG cells’ pairwise responses (Figure S2C). Most pairwise interactions were saturated additive (gray), but some showed antagonism (red) or suppression (black) (Figures 3C, S2C-F). Combining the full set of pairwise interactions with measurements of individual ligand strength enabled clustering of ligands based on their functional similarity (Figure 3C, dendrogram). Ligands appearing in the same cluster (group) exhibit the same pattern of interactions with other ligands, suggesting they function interchangeably across pairwise ligand combinations. These groups can be represented compactly around a circle (Figure 3D), with the grayscale level of each group indicating the approximate individual activity of its ligands and internal lines representing the non-additive interactions between groups. These groups included [BMP2, BMP4], [BMP5, BMP6, BMP7, BMP9], [BMP10], [GDF5, GDF7], and [GDF6]. Here and below, square brackets denote sets of ligands that constitute an equivalence group.

In NMuMG cells, the full set of interactions among 10 ligands reduces to a set of five non-additive interactions among five distinct ligand groups. Ligands in the first group, [BMP5, BMP6, BMP7, BMP9], activated strongly as individuals and interacted additively with one another and all other ligands, except for weak antagonism with BMP10. The second group, [BMP2, BMP4], also contained strong activators, but was distinguished by its susceptibility to antagonism by GDF5 and GDF7 and suppression by BMP10. The third and fourth groups, [GDF5, GDF7] and [GDF6], comprised non-activating ligands that respectively did or did not antagonize [BMP2, BMP4]. Finally, [BMP10] formed its own equivalence group based on its unique interactions and intermediate strength. Thus, most of the ligand distinctions represented here emerge from the interactions and could not have been inferred from signaling properties of the individual ligands.

This systematic, interaction-based classification of ligands provides a multi-ligand perspective for predicting the effects of ligand combinations expressed in biological processes. For example, BMP2, BMP4, and BMP7 are coexpressed in skeletal development, but different combinatorial knockouts generate distinct skeletal defects (Bandyopadhyay et al. 2006). These results implied a unique role for BMP7, consistent with its classification in a separate equivalence group from BMP2 and BMP4. However, the equivalence map determined in NMuMG cells is not necessarily universal to all cell types. BMP receptor expression is variable across cell types in developing tissues (Danesh et al. 2009), and changes in receptor expression can significantly alter pairwise ligand interactions (Antebi et al. 2017). Therefore, these results provoke the question of how equivalence groups vary with cell context in general and with receptor expression profiles more specifically.

### Ligand equivalence depends on cell context

Mouse embryonic stem cells (mESCs) present an ideal cell context in which to study BMP signaling. Manipulation of BMP signaling in mESCs is a key step in many directed differentiation protocols (Craft et al. 2013; Gouon-Evans et al. 2006), and single BMPs and pairs can produce distinct cell fates in the same protocol (Andrews et al. 2017). mESCs also differ from NMuMG cells in their expression of three of the five BMP receptors, providing a different BMP receptor context (Figure 4A). Finally, the two cell types are known to respond in distinct ways to some ligand combinations (Antebi et al. 2017).

**Figure 4:**
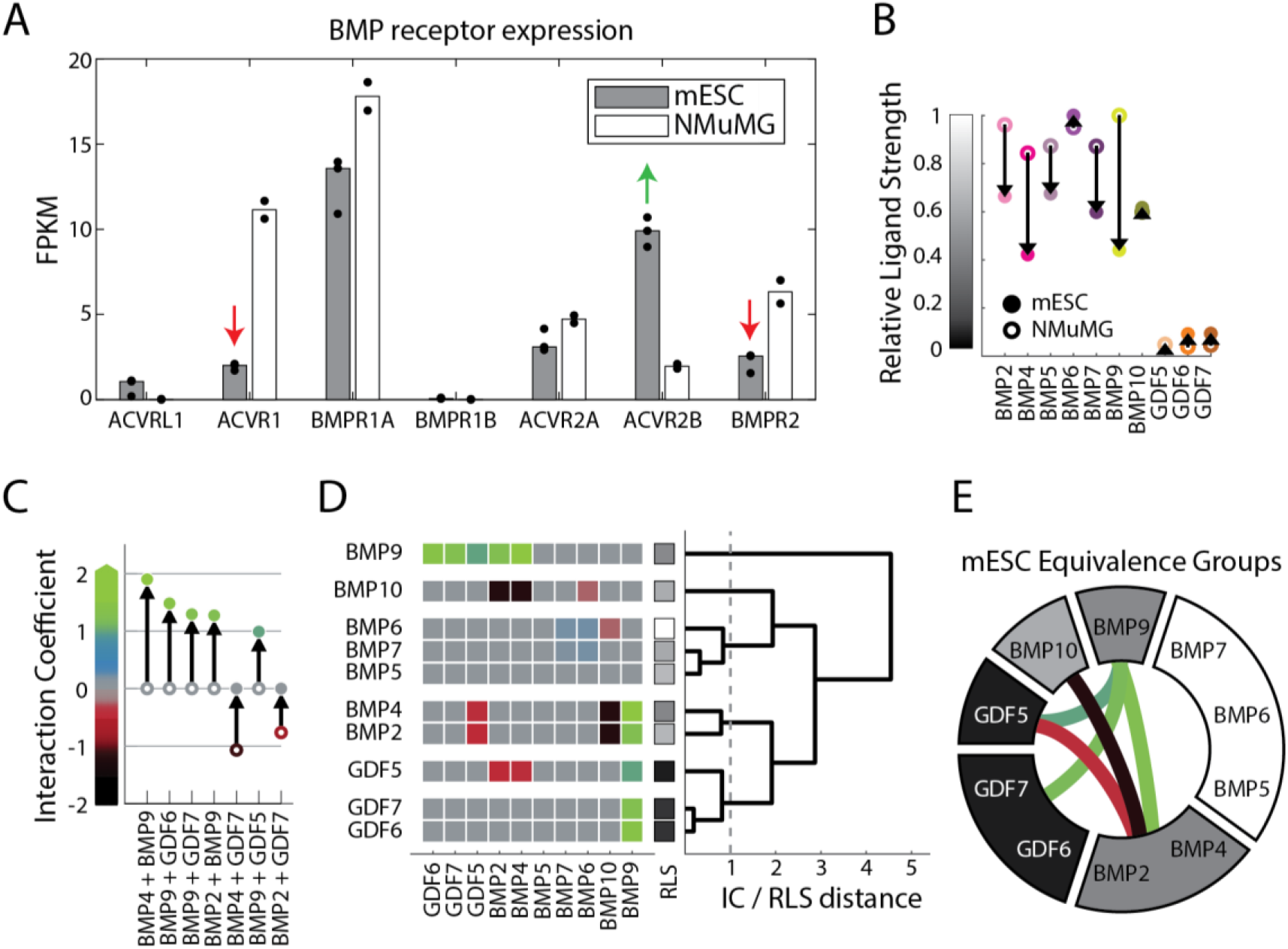
Mouse ES cells exhibit distinct BMP equivalence groups. (A) RNA-Seq of BMP receptors in mESC reporter and wild-type NMuMG cells show differences in receptor expression (highlighted by green and red arrows) between the two cell lines, with mESCs expressing more ACVR2B, but less ACVR1 and BMPR2. Dots show at least two biological repeats, and bar height is median. (B) RLS values in mESCs (closed circle) differ from NMuMG cells (open circle, cf. Figure S1G), with more intermediate strength activators in mESCs. Arrows indicate the change in mESCs relative to NMuMG cells. (C) IC values for seven pairwise interactions changed significantly (|ΔIC|>0.5) between NMuMG cells (open circle) and mESCs (closed circle). The majority show a net increase of IC, producing five new synergistic interactions. Both the color and y-axis value of the open and closed circles indicate the IC value for the indicated cell lines, computed on the median of four biological repeats. (D) Hierarchical clustering of IC and RLS values in mESCs reveal a new set of ligand equivalence groups compared to NMuMG cells (cf. Figure 3D), using the same cutoff criteria to group ligands by similar individual and pairwise interactions (STAR Methods). The square colors indicate either the ligand’s IC value when paired with another ligand or the RLS grayscale for its individual strength (see y-axis colormaps in B and C), as indicated by x-tick labels. The dendrogram on the right-hand side shows complete-linkage clustering of Euclidean distance between ligand features. (E) BMP equivalence in mESCs, as classified by hierarchical clustering in D, includes six equivalence groups. Grayscale value indicates the approximate RLS of the grouped ligands, while connecting line colors indicate the pairwise interactions between groups of ligands. Groups without a linking edge interact additively. See also Figure S3.

We used a fluorescent mESC reporter line to analyze the dose response for each of the ten ligands (STAR Methods). mESCs showed strikingly different responses to the individual ligands compared to NMuMG cells. They exhibited a greater diversity of activation strengths (Figure 4B), as well as greater EC_50_ values and lower Hill coefficients, indicating larger input dynamic ranges with lower sensitivity to moderate ligand concentrations (Figure S3A). In particular, high concentrations (approaching 10 μg/mL) of some ligands failed to fully saturate the response.

Ligand interactions also differed between the two cell lines. While most pairwise interactions remained saturated additive (Figures S3B-D), a subset showed marked differences (|ΔIC|>0.5) between the two cell lines (Figure 4C). Most notably, BMP9 acquired synergistic interactions with activating ligands [BMP2, BMP4] as well as with non-activating ligands [GDF5, GDF6, GDF7] (Figure S3E), an effect proposed to emerge when ligands in pairs, but not individually, exclusively form high efficiency complexes, due to competition for a limited receptor pool (Antebi et al. 2017). At the same time, strong antagonistic interactions between GDF ligands and [BMP2, BMP4] were lost. Together, these shifts represented an overall decrease in antagonism and increase in synergy across all ligand pairs (Figure 4C). Combined with the changes in individual activation, these differences generated new ligand equivalence relationships (Figure 4D). While some groups and interactions were preserved, such as the suppression between [BMP2, BMP4] and [BMP10], many other groups were split or merged (cf. Figures 4E, 3D).

Together, these results directly demonstrate the contextuality of ligand interactions and ligand equivalence relationships. Within this larger set of pairwise interactions, a randomly chosen pair of ligands is most likely to interact additively, but is nevertheless unlikely to function equivalently in terms of its individual potency and pairwise interactions with other ligands. Moreover, equivalence groups can change in new cell types. Therefore, an observation of equivalence in one cell context does not guarantee equivalence in another, and vice versa.

### ACVR1 knockdown recapitulates features of mESC ligand interactions

A key factor that could explain differences in ligand equivalence between NMuMG cells and mESCs is receptor expression (Danesh et al. 2009; Panchision et al. 2001; Yamauchi, Phan, and Butler 2008). mESCs express less ACVR1 and BMPR2, but more ACVR2B, than NMuMG cells (Figure 4A). Out of these three receptors, we first focused on ACVR1, a Type I receptor that plays key roles in bone and brain diseases (Shore et al. 2006; Valer et al. 2019; Agarwal et al. 2017), and asked whether reducing its expression could make ligand responses in NMuMG cells more similar to those of mESCs. To achieve stable knockdown of ACVR1, we constitutively expressed shRNA against ACVR1 in the NMuMG reporter cell line. This perturbation reduced expression to less than 20% of wild-type levels with minimal effects on off-target receptors (Figure S4A) and had the same effects on ligand interactions as transient siRNA knockdown of ACVR1 (Figure S4B).

ACVR1 knockdown recapitulated some of the observed differences between NMuMG cells and mESCs, particularly those involving BMP9, a ligand known to signal through ACVR1 (Valer et al. 2019). BMP9, the strongest individual activator of NMuMG cells, signaled only weakly in the knockdown cells, consistent with its low activity in mESCs (Figure 5A, cf. 4B). More dramatically, ACVR1 knockdown reproduced the synergy of BMP9 with GDF5, GDF6, and GDF7 observed in mESCs (Figures 5B, S4G,H, cf. 4C). Finally, in both mESCs and ACVR1 knockdown, the equivalence of BMP9 with BMP5, BMP6, and BMP7 was eliminated (Figures 5C,D, cf. 4E).

**Figure 5:**
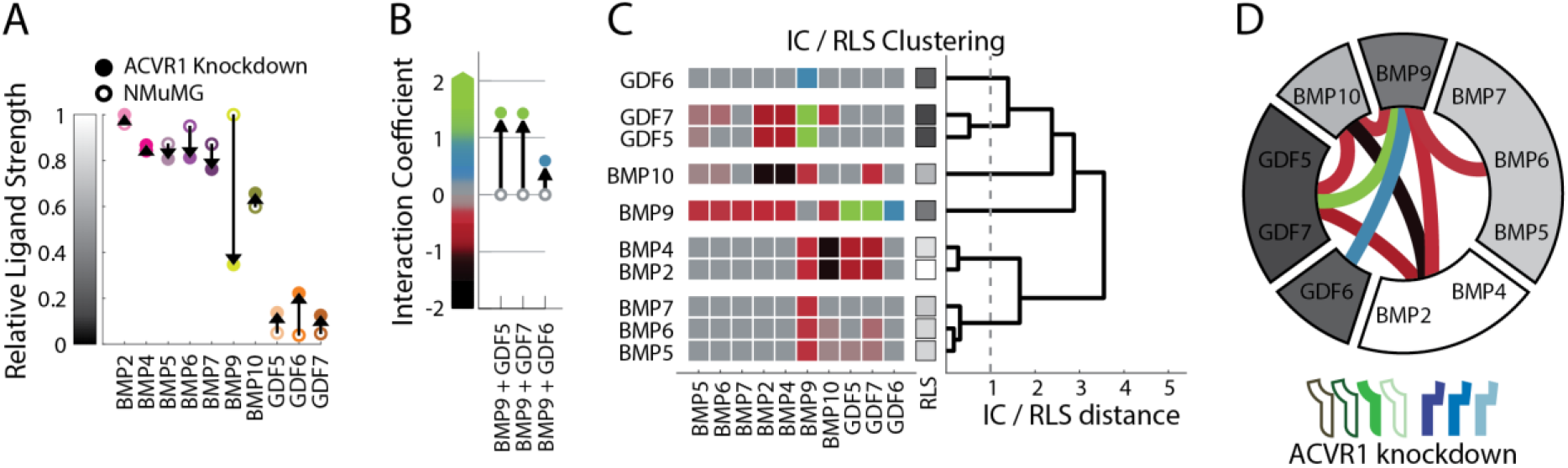
ACVR1 knockdown reveals the effects of receptor context on ligand equivalence. (A) ACVR1 knockdown (closed circle) alters RLS values compared to NMuMG cells (open circle, cf. Figure S1G). BMP9 is the most strongly affected. (B) Three pairwise interactions changed significantly (|ΔIC|>0.5) between NMuMG cells (open circle) and ACVR1 knockdown (closed circle), producing new synergistic interactions with BMP9. Open and closed circles are colored by IC value. (C) Hierarchical clustering of RLS and IC values (as in A and B) in the ACVR1 knockdown context shows a more complex equivalence map. The square colors indicate either the ligand’s IC value when paired with another ligand or the RLS grayscale for its individual strength (see y-axis colormaps in A and B), as indicated by x-tick labels. The dendrogram on the right-hand side shows complete-linkage clustering of Euclidean distance between ligand features. (D) BMP equivalence following ACVR1 knockdown, as classified by hierarchical clustering in C, includes six groups. Wedge color shows approximate RLS value of the contained ligands, while connecting line colors indicate pairwise interactions between groups of ligands. Groups without a linking edge interact additively. Here, as in panels E-H, each receptor perturbation is shown as a cartoon of filled and empty receptors, which are drawn in the following order: ACVRL1, ACVR1, BMPR1A, BMPR1B, ACV2A, ACVR2B, BMPR2. See also Figures S4.

By contrast, other distinct features of the mESC response did not appear in the ACVR1 knockdown. Certain non-additive interactions in NMuMG cells persisted even after ACVR1 knockdown (Figures 5C, S4C-E). For example, [BMP2, BMP4] was still suppressed by [BMP10] (Figures 5C, S4F) and antagonized by GDF5 (Figure 5C). Evidently, significant expression of ACVR1 is not necessary for these non-additive interactions. ACVR1 knockdown also produced new interactions not observed in NMuMG cells or mESCs. [BMP9] antagonized, rather than synergizing with, the other strongly activating ligands, including [BMP2, BMP4] and [BMP5, BMP6, BMP7]. This result therefore also showed that a single ligand (BMP9) can produce interactions of opposite signs in the same cell context, which was also not observed in either NMuMG cells or mESCs. Together, these results suggest that differences in receptor expression, and ACVR1 expression more specifically, can account for some of the distinct ligand responses of mESCs and NMuMG cells, and reveal that knockdown of a single receptor subunit can substantially increase the complexity of ligand interactions.

### Perturbations of multiple BMP receptors reveal flexibility of ligand equivalence

To further explore the effects of single receptor perturbations on ligand equivalence, we individually knocked down the two most abundant receptors, BMPR1A and BMPR2, or ectopically expressed the two receptors with weakest endogenous expression, ACVRL1 and BMPR1B. These four receptors differ widely in their ligand selectivity and expression profiles. BMPR2 and BMPR1A bind a diverse set of BMPs (Ten Dijke et al. 1994; Nishitoh et al. 1996; Orriols, Gomez-Puerto, and Ten Dijke 2017) while ACVRL1 and BMPR1B have more specific ligand preferences. Additionally, BMPR1A and BMPR2 are broadly expressed across distinct tissues, whereas BMPR1B and ACVRL1 are highly expressed in relatively few tissues (Danesh et al. 2009; Cunha and Pietras 2011). We reasoned that perturbing receptors with such diverse properties could provide insight into how different specificities of receptor-ligand interactions influence ligand equivalence.

To explore these diverse receptor properties, we first generated stable shRNA knockdown cell lines for BMPR1A and BMPR2, similar to the ACVR1 line (Figures S5A,B). Knockdown of BMPR2, which is also reduced in mESCs relative to NMuMG cells, decreased activation by BMP2 and BMP4, removed their antagonism by GDF5 and GDF7, and fused [GDF5, GDF6, GDF7] into one equivalence group (Figures 6A “BMPR2 KD,” S5C-G), all of which were features of BMP signaling in mESCs. Thus, this perturbation recapitulated certains aspects of mESC equivalence, though distinct from those produced by ACVR1 knockdown. Knockdown of BMPR1A produced a further simplified version of the BMPR2 knockdown, with no activation by BMP2 and BMP4 and only one non-additive interaction occurring between two of the three groups (Figures 6A “BMPR1A KD,” S5H-J). The loss of most non-additive interactions following knockdown of BMPR2 or BMPR1A also revealed that these receptors, unlike ACVR1, are necessary for many non-additive ligand interactions in NMuMG cells.

**Figure 6:**
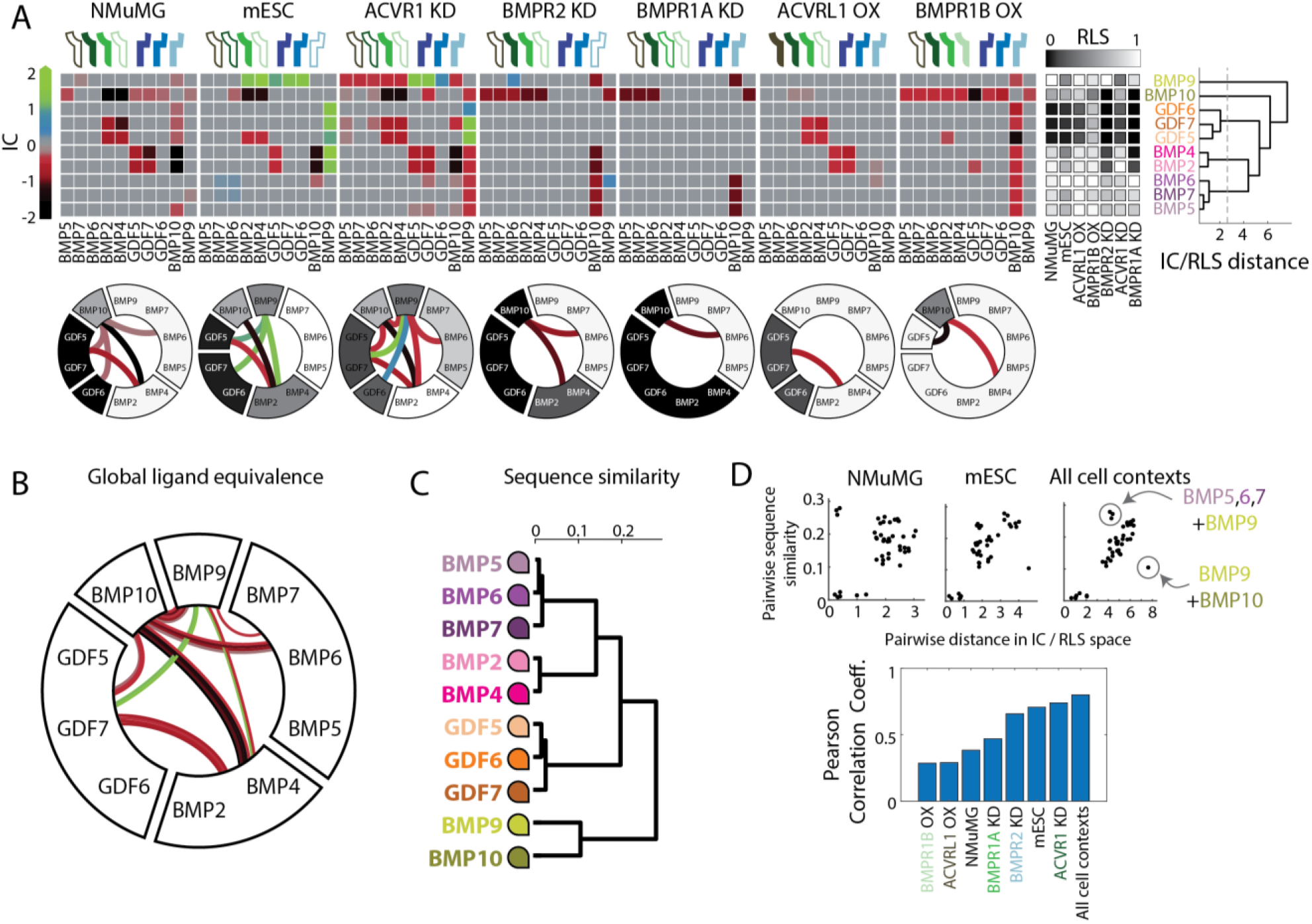
Analysis of multiple cell contexts reveals global ligand equivalence groups. (A) Clustering RLS and IC values for all ligands and pairs across all cell contexts shows global ligand similarity (upper row). The full set of IC values show the distribution of ligand interaction types (colors based on Figure 3C) and identify ligands with different frequencies of non-additive interactions. Lower row shows the corresponding ligand equivalence groups (reproduced from Figures 3D, 4E, 5D-H). For the dendrogram (right), clustering was done by complete linkage of the Euclidean distance between RLS and BC features, as before. (B) The global equivalence map was determined by cutting the dendrogram (panel A, right) at the indicated distance. Internal links show all instances of non-additive interactions between those ligands. Global groups tend to have linkages of the same interaction type (i.e. similar colors), with widely varying frequency. For example, BMP2 and BMP4 almost always interact with BMP10 but never with BMP5. (C) Recombinant ligand sequences were clustered by complete-linkage clustering of pairwise alignment distance, as computed by the BLOSUM50 matrix between globally-aligned sequences, neglecting all gaps. Ligands comprise roughly four groups of related proteins, indicated by color families. The recombinant ligands used in this study constitute the mature secreted form of the ligand, exclude pro-domains, and vary in length from 108 to 146 amino acids (see also Figure S7A). (D) Sequence similarity, quantified as in (C), correlates with IC/RLS similarity observed in each cell context, though the extent of correlation varies. In the correlation across all cell contexts, notable outliers include BMP9’s surprising functional difference from BMP10, despite their significant sequence similarity, and BMP9’s unexpected similarity with BMP5, BMP6, BMP7, despite relatively dissimilar sequences. Pearson correlation coefficient (bottom) quantifies correlation between sequence similarity and IC/RLS distance for each cell line. Highest correlation is achieved by considering all cell contexts together. See also Figure S5, S6, and S7.

Thus far, perturbations were of relatively promiscuous receptors, known to interact with a broad spectrum of ligands. By contrast, the receptors ACVRL1 and BMPR1B are known to be more ligand-specific (Chai et al. 2020; Nishitoh et al. 1996; Cunha and Pietras 2011), primarily binding BMP9 and BMP10 or the GDFs, respectively. We therefore ectopically expressed these receptors by stably integrating constructs encoding receptor cDNA into the NMuMG reporter cell line (Figure S6A). Ectopic ACVRL1 expression increased activation by BMP10 and removed its many suppressive and antagonistic interactions with other ligands (Figure 6A “ACVRL1 OX,” S6B-D). Similarly, BMPR1B increased activation by its preferred ligands GDF5, GDF6, GDF7, and removed their antagonism of BMP2 and BMP4 (Figure 6A “BMPR1B OX,” S6E-H). Thus, both perturbations reduced the number of equivalence groups and non-additive interactions, with ectopic BMPR1B expression allowing 8 of the 10 ligands to signal equivalently. More specifically, ectopic expression of each receptor resulted in its preferred ligands becoming equivalent to other strongly activating, weakly interacting ligands, such as BMP5, BMP6, BMP7, and BMP9. These experiments suggest that non-additive interactions could arise from receptor competition, which is specified by the receptor’s ligand preferences and can be relieved by ectopic receptor expression.

### Comparing signaling across cell types reveals global structure of ligand equivalence

Together, the pairwise interactions and corresponding equivalence maps observed across all cell contexts provide a more global view of the structure of BMP signaling. The full set of pairwise interactions reveals the striking plasticity of ligand strength and interactions (Figure 6A, top). GDF5, GDF6, and GDF7 were usually weak activators, while BMP5, BMP6, and BMP7 strongly activated all cell types. The remaining ligands were active in some but not all receptor contexts. While most pairwise interactions were additive, every ligand participated in synergy or antagonism with another ligand in at least one receptor context. Non-additive interactions tended to be consistent for a given ligand pair, exhibiting the same interaction type (i.e. color) across cell lines. All synergistic ligand interactions involved BMP9, whereas all suppressive interactions involved BMP10. Considered as a system, the meaning of any given BMP ligand is thus inherently contextual, depending both on other ligands and on receptor context.

To represent the full dataset in a more compact way, we clustered ligands according to the similarity of their interactions across receptor contexts (Figure 6A, right), such that ligands in the same cluster had the same individual strength and pairwise interactions across all receptor contexts. This approach classified the ten BMPs into five global equivalence groups: [BMP2, BMP4], [BMP5, BMP6, BMP7], [BMP9], [BMP10], and [GDF5, GDF6, GDF7]. [BMP2, BMP4], as well as [BMP5, BMP6, BMP7], exhibited the strongest version of equivalence. Despite frequent changes in individual or combinatorial signaling across cell contexts, responses to ligands within each group always changed in the same manner, such that the ligands in each group were equivalent in all contexts. The global equivalence map also illustrates the variability and prevalence of ligand interactions (Figure 6B). For example, interactions between [BMP9] and [BMP2, BMP4] can vary from synergy to antagonism. In all receptor contexts except one, [BMP10] interacts non-additively with [BMP2, BMP4], while [BMP5, BMP6, BMP7] have only additive interactions with [BMP2, BMP4] or [GDF5, GDF6, GDF7]. (Note that we grouped GDF6 with GDF5 and GDF7 despite its apparent lack of antagonistic interaction with [BMP2, BMP4] because its higher overall EC_50_ may obscure a potential underlying antagonism (Figure S6E, inset; STAR Methods).)

This global equivalence diagram also portrays another surprising feature of context-dependent ligand differences. A single linear ligand ordering, following the circle from BMP9 around to BMP10, is consistent with all observed equivalence maps. With this ordering, no rearrangements of ligand position are necessary in any observed context. This suggests that ligand differences fall on a linear continuum, and is distinct from what one would expect in a clustering-based analysis, where many orderings are consistent with the same linkage tree. While analysis of additional receptor contexts could further increase the number of global equivalence groups, this diagram efficiently summarizes ligand differences and interactions.

How closely do the equivalence relationships assessed here reflect intrinsic aspects of ligand sequence? To assess this, we compared similarity of ligand protein sequence (Figures 6C, S7A) with similarity of pairwise interaction profiles. In individual cell lines, these two metrics were only partially correlated (Figure 6D, top), suggesting that ligand sequence similarity does not fully predict functional similarity in a single receptor context. For example, BMP5 and BMP9 act equivalently in NMuMG (Figure 3D) but are relatively dissimilar in sequence (Figure 6C). Ligand sequence correlated with ligand function more strongly when all cell lines were considered simultaneously (Figure 6D, bottom). This ‘global’ correlation coefficient, 0.80, approached the maximum value obtained with randomly sampled cell line combinations, 0.84 (Figure S7B). Interestingly, however, even when all cell lines were considered together, BMP9 and BMP10 exhibited more divergent functional behavior than expected given their sequence similarity. Overall, this analysis suggests that ligand sequence similarity does not generally reflect functional similarity in any individual cell context, even for the most highly homologous ligands. Instead, it appears to correlate more strongly with aggregated, or global, functional similarity across multiple contexts.

### A mathematical model of competitive ligand-receptor interactions can explain context-dependent ligand interactions

As shown above, ligands can interact in different ways depending on receptor context. To understand how such complex, contextual responses could emerge, we analyzed a mathematical model of the pathway developed in a parallel study (Su et al. 2020). This model assumes that each ligand variant, *L_i_*, can bind to any pair of Type I and Type II receptors, *A_j_* and *B_k_* respectively, with affinity *K_ijk_* to produce a trimeric, ligand-receptor signaling complex, *T_ijk_* (Figure 7A), whose binding is captured by a set of ordinary differential equations describing mass-action kinetics. Each signaling complex can then phosphorylate SMAD proteins with its own specific kinase activity *ε_ijk_*. Receptors are expressed at specific total levels, 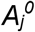 or 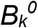. The limited total amount of each receptor generates competition among ligands for available receptors. Other features of the BMP system—including step-wise assembly of ligand-receptor complexes, the heterotetrameric stoichiometry of actual receptor complexes, co-receptors, and other factors—play important roles in the natural system but were not required to explain observations in this work (STAR Methods).

**Figure 7:**
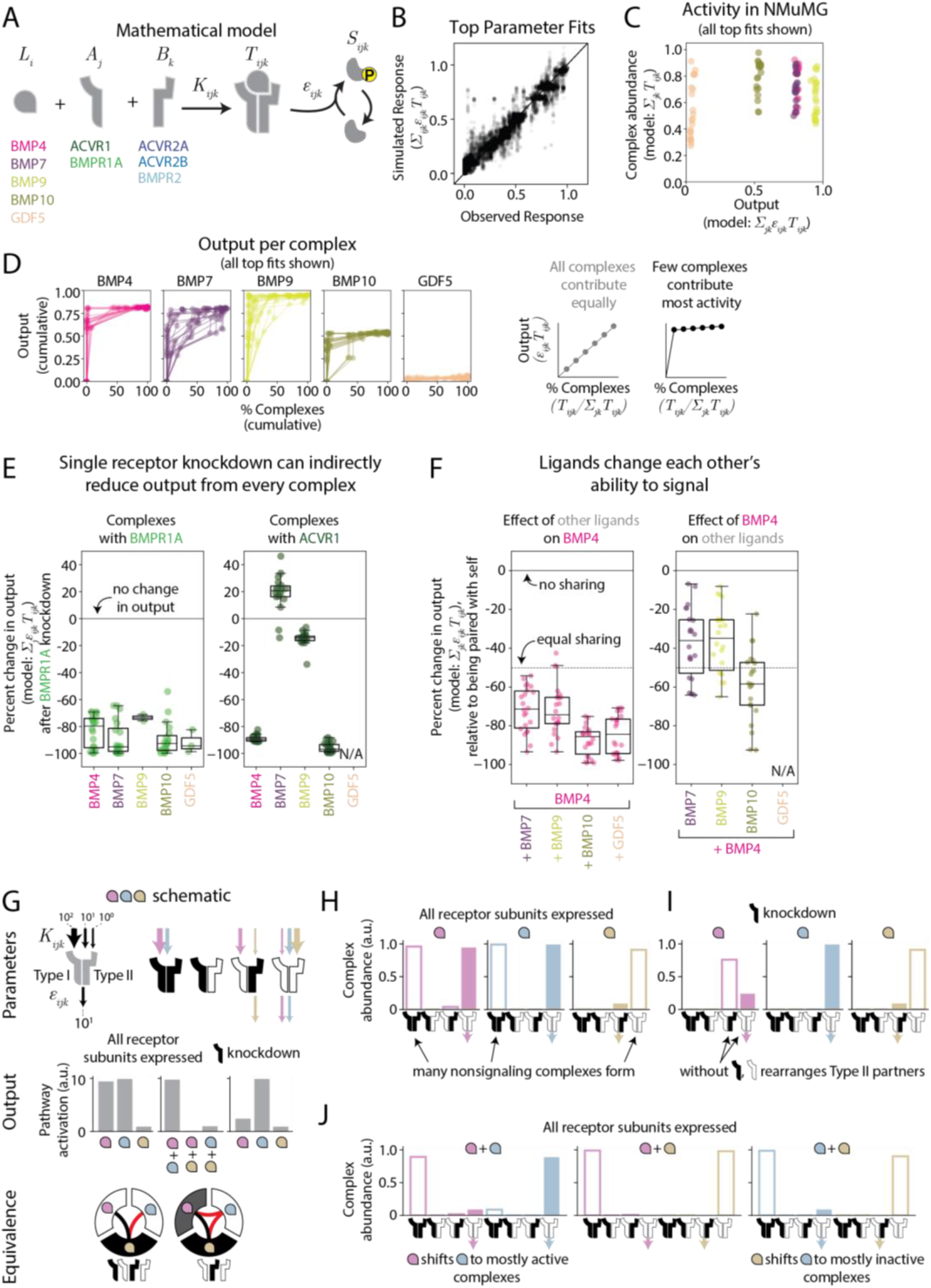
Mathematical model of receptor competition can explain contextual ligand equivalence groups. (A) In a minimal BMP receptor competition model, mass action kinetics govern one-step assembly of ligand (*L_i_*), Type I receptor (*A_j_*), and Type II receptor (*B_k_*) into trimeric signaling complexes (*T_ijk_*) that activate pathway output with some activity (*ε_ijk_*). The included components are listed beneath their associated variable. (B) Observed responses, normalized between 0 and 1, correlate with simulated responses in top parameter fits. (C) Total complex abundance (in arbitrary units, normalized to total possible number of complexes) does not correlate with overall output from a given ligand. Ligands of intermediate strength (e.g. BMP10) can bind more receptors than strongly activating ligands. (D) For individual ligands activating NMuMG, the inferred output of each complex is plotted against its percentage of the total complexes for all top parameter fits. A minority of the complexes that form can produce the majority of the output. (E) The percent change in output (relative to NMuMG) from BMPR1A- and ACVR1-containing complexes following BMPR1A knockdown is shown for all five ligands. BMPR1A knockdown reduces the number of BMPR1A-containing complexes and reduces their output as expected, but also can reduce output from ACVR1-containing complexes, as BMPR1A loss increases availability of Type II receptors. All top parameter fits are plotted, unless the absolute change is small (i.e. <0.05 change in response). GDF5, which does not activate these cells and has only small changes in ACVR1 output, is not shown (N/A). (F) Percent change in output (relative to signaling alone) from complexes containing BMP4 or another ligand are shown for BMP4 paired with four other ligands: BMP7, BMP9, BMP10, and GDF5. Equal sharing between the two ligands would reduce output by 50%, whereas ligands that do not share complexes produce as much as output alone as they do in a pair. All top parameter fits are plotted, unless the absolute change is small (i.e. <0.05 change in response). GDF5, which does not activate these cells and has only small changes in overall output, is not shown (N/A). (G) A schematic parameter set shows three ligands with different affinities and activities for four possible receptor dimers. In the model, these ligands have different individual strengths and pairwise strengths which change with receptor perturbations. (H) In a simulation where each receptor subunit is available, the three ligands form both signaling and nonsignaling complexes. Bar height shows complex abundance. Empty bars show nonsignaling complexes, whereas filled bars show signaling complexes. (I) After simulated knockout of the black Type I receptor, the abundances of many signaling complexes change, even if they did not involve the knocked out receptor. (J) In a simulation where each receptor subunit is available, competition for shared receptors changes the formation of signaling complexes by each ligand. The pink and gold ligands can respectively shift the blue ligand to preferentially form its signaling and nonsignaling complexes. See also Figure S8.

To identify parameter values that can generate the range of interactions observed experimentally, we focused our analysis on a single ligand from each global equivalence group. Similarly, we considered only the five broadly-expressed receptors (Figures 7A, S4A, S5A): the Type I receptors ACVR1 and BMPR1A and the Type II receptors ACVR2A, ACVR2B, and BMPR2. We fit the model simultaneously to measurements of individual and pairwise responses in the four NMuMG-derived cell lines whose receptor profiles are restricted to this set (i.e. NMuMG and the knockdowns of ACVR1, BMPR2, and BMPR1A). We then analyzed a set of 22 solutions from nearly 7,000 least-squares fits that produced the lowest fitting error and preserved key pairwise interactions (Figures 7B, S8A; STAR Methods).

Within these best fit parameter sets, all ligands were promiscuous, with substantial but variable affinities for most or all receptors (Figure S8B, top). BMP7 and GDF5 had lower affinities overall across all complexes, while BMP4, BMP9, and BMP10 had higher affinities for select complexes (Figure S8B, top). The ligands also differed in the activity of the resulting complexes. GDF5 weakly activated most signaling complexes, while other ligands strongly activated multiple receptor complexes (Figure S8B, bottom). Many signaling complexes exhibited strong affinity but weak activity or vice versa (Figure S8C). A parallel computational study of the BMP pathway showed that this inverse relationship between activity and affinity generates the complex multi-ligand responses necessary for combinatorial addressing, in which ligand combinations can selectively activate cell types (Su et al. 2020). Taken together, these results suggest that the context-dependent ligand interactions observed experimentally can be explained by affinity-based competition to form complexes with different activities. They further suggest that the effective biochemical parameters for mammalian BMPs exhibit the features associated with combinatorial addressing.

### In the model, context-dependence emerges from redistribution of signaling complexes

We identified three key features of the best fit parameter sets that allowed them to reproduce the observed context-dependent ligand interactions. First, each of these parameter sets generated many high affinity (high *K_ijk_*), low activity (low *ε_ijk_*) signaling complexes. Ligands can appear weak not because they form fewer complexes but rather because they preferentially form complexes with weaker activity (Figure 7C). More surprisingly, even for highly active ligands, most of the response was generated by a small fraction of the formed complexes (Figures 7D, S8E,F).

A second feature of top parameter fits was that they allowed individual receptor subunits to participate in multiple complexes with widely varying activities. Consequently, when receptors are rearranged into new complexes following the perturbation of a single receptor, significant changes in pathway activity can result. These changes can be direct and indirect. For example, knockdown of a single receptor subunit can directly reduce the output from complexes in which it appears by reducing their concentrations. It can also indirectly affect the response from other signaling complexes by increasing or decreasing abundance of inactive, competing complexes (Figures 7E, S8G,H). For example, in the model, knockdown of BMPR1A reduces the activity of BMP4 and BMP10 complexes containing ACVR1 (and lacking BMPR1A itself). This indirect effect is due to the release of ACVR2B from BMPR1A, which allows formation of ACVR1-ACVR2B complexes that compete with the higher-activity ACVR1 complexes (Figures S8G,H). These indirect effects account for much of the complexity of receptor context.

Finally, the parameter fits also produced hidden, or compensatory, shifts in the activity of one ligand in response to addition of a second ligand. For example, in NMuMG cells, BMP4 combined additively with BMP7 (Figure 3D), seemingly implying that the two ligands do not affect each other’s ability to signal. However, in nearly all top fitting results, BMP7 produced more output than expected if it shared its receptors equally with BMP4, while BMP4 produced less than expected. Thus, the saturated additive BMP4-BMP7 interaction resulted from opposite effects on the underlying signaling activities of the two ligands (Figures 7F, S8I,J). A more complete analysis of redistribution of signaling complexes is shown in Figures S8D-J, which highlight a single random parameter set.Taken together, these results show how reassortment of ligand and receptor subunits can reproduce observed interactions through sometimes non-intuitive mechanisms.

To better illustrate how these effects occur, we constructed a simplified hypothetical system with three ligands, two Type I, and Type II receptor subunit variants interacting with the affinity and activity parameters shown in Figure 7G. As observed in the parameter fits, when all receptor subunits are expressed, even the highly active pink and blue ligands form many non-signaling complexes (Figure 7H). Further, knocking down a receptor indirectly reduces output from unperturbed receptors (Figure 7I). For example, when the black Type I receptor is removed, the white Type I receptor obtains unrestricted access to the pool of Type II receptors, allowing formation of a new, lower-activity complex with the pink ligand. Thus, even though the white Type I receptor is not itself perturbed, its output is reduced.

This schematic example also shows how addition of one ligand can modify the activity of another through competition. For example, the gold ligand competes with the blue ligand to bind to the white Type I and II receptor, which is the only receptor through which the blue ligand can signal (Figure 7J), producing an antagonistic interaction between the two ligands. By contrast, when the blue and pink ligands are combined, signaling by the former is increased while signaling by the latter is decreased relative to their individual activities. These changes cancel each other out, resulting in minimal change to the total signal. In this case, the apparent additivity of the interaction conceals compensatory changes in the distributions of both ligands. Thus, competition generally redistributes ligands across receptor complexes but does not necessarily produce changes in total signaling activity. This simplified example illustrates many of the principles observed in the full model.

## Discussion

Individual BMP ligands are known to preferentially bind and signal through specific receptors. However, preferences alone cannot explain the striking contextuality of BMP ligand responses. Here, we introduced contextual ligand equivalence groups as an alternative framework to understand ligand differences when signaling in combinations across receptor contexts. This approach uses systematic pairwise measurements to define equivalence groups, each comprising a set of ligands that interact similarly with all other ligands when activating a given cell type (Figure 3D). Critically, these equivalence groups are contextual, changing with alterations in receptor expression (Figures 4–6). In contrast to ligand classifications based on biochemical properties like affinity, contextual equivalence groups classify BMP ligands and receptors based on their emergent functional properties.

Contextual equivalence groups can explain ligand redundancy that varies between developmental contexts. For example, BMP10 signals in both developing heart and vasculature (Chen et al. 2013). Although BMP9 has the closest sequence similarity to BMP10, the contextual equivalence maps show that BMP9 can only substitute for BMP10 in the presence of ACVRL1 (Figure 5G). Consistent with this picture, BMP9 rescued loss of BMP10 in developing mouse vasculature but not heart, as ACVRL1 expression is limited to vascular epithelium (Chen et al. 2013; Seki, Yun, and Oh 2003). Contextual equivalence groups also reveal the effects of ligand context. For example, in receptor contexts where BMP4 and BMP7 are strong activators, they can only replace each other when antagonizing ligands such as GDF5, GDF6, GDF7, or BMP10 are absent (Figure 3D, 5G, 5H). Consistent with this, BMP4 and BMP7 functioned redundantly in the developing kidney (Oxburgh et al. 2005), where those antagonizing ligands are not expressed (Godin, Robertson, and Dudley 1999). By contrast, loss of BMP4 produced more pronounced effects than loss of BMP7 in the context of developing skeleton with conditional BMP2 knockout (Bandyopadhyay et al. 2006), a context where GDF5 is expressed and plays a key role (Storm and Kingsley 1999).

The contextual equivalence analysis of BMP signaling introduced here can be improved and extended in several ways. First, analyzing ligand equivalence in additional receptor contexts would provide more complete coverage of possible BMP signaling contexts. Second, including heterodimeric ligands such as BMP2/7 could reveal whether these ligands have distinct contextual properties from their related homodimers, in addition to their unique individual properties (Valera et al. 2010). Third, many natural contexts use three or more ligands simultaneously. The approach taken here could be extended to combinations of three or more ligands to determine whether pairwise interactions completely determine multi-ligand effects or whether higher-order interactions are important.

Contextual equivalence groups can be explained by a simple biochemical model, suggesting that the context-dependent behavior of BMP ligands could arise from differences in receptor affinity and signaling complex activity. Key aspects of this model are consistent with previously observed features of BMP signaling. For example, loss of BMP or related TGF-β receptors indirectly affects the abundance of other receptors complexes (Hiepen et al. 2019; Lowery et al. 2015; Razavi, Lopez, and Lyons 2019). More specifically, BMPR2 knockdown has unexpected and ligand-specific effects across a variety of cell types (Han et al. 2013; Hiepen et al. 2019; Yu et al. 2005; Olsen et al. 2018; Theilmann et al. 2020), which may be explained by the low predicted activity of BMPR1A-BMPR2 complexes that recurred in all model fits. In pulmonary artery cells and myeloma cells, BMPR2 knockdown either did not affect or decreased signaling by BMP4 and BMP10, while, respectively, increasing or having no effect on signaling by BMP7 and BMP9. In the model, without BMPR2, BMP4 and BMP10 preferentially formed lower activity complexes than with BMPR2. Therefore, the absence of BMPR2 redistributes receptors to reduce pathway activity overall. By contrast, BMP7 and BMP9 mostly form complexes with high activity in the absence of BMPR2. For these ligands, association with other Type II receptors would tend to increase or maintain pathway activation. Thus, while the minimal phenomenological model established here does not represent the full complexity of the BMP system, it provides a foundation for a unified, systems-level framework for describing ligand differences, generating hypotheses, and reconciling otherwise counter-intuitive effects of perturbations among cell types.

Independent experimental measurements of the effective affinity and activity parameters for various ligand-receptor signaling complexes (Figure 7) could determine parameters not fully constrained by existing data. This would allow the model to predict responses to ligand combinations in arbitrary cell (i.e. receptor) contexts. More specifically, measuring the effective affinity of overall complex formation may be a crucial follow-up to measuring the affinity of BMP ligands binding single receptors (Heinecke et al. 2009; Saremba et al. 2008; Aykul and Martinez-Hackert 2016; Isaacs et al. 2010), and measurements of signaling complex activity may provide more examples of BMP ligands that bind and only weakly activate receptors, or vice versa (Nishitoh et al. 1996; Nickel and Mueller 2019; Aykul et al. 2020). Finally, while the existing model shows that competitive ligand-receptor interactions are sufficient to explain key features of contextual equivalence groups, further tests will be essential to understand how additional biochemical interactions, such as binding to negative regulators like Noggin, could enable other pathway behaviors.

Ligand classification by pairwise interactions can generalize beyond the core BMP pathway. First, additional TGF-β family ligands, such as TGF-β or activin, could be included. These ligands regulate phosphorylation of the related SMAD2/3 transcription factors, share some but not all receptors with BMP ligands (Razavi, Lopez, and Lyons 2019), and exhibit distinct activation dynamics that may allow more complex ligand integration (Weiss and Attisano 2013; Zi, Chapnick, and Liu 2012). Expanding into this related set of ligands not only introduces the probability of more complex equivalence relationships but also provides deeper insight into a critical signaling system in cancer and wound healing (Bierie and Moses 2010; Pakyari et al. 2013). Second, analyzing a broader set of BMP targets, including non-canonical outputs and broad repertoires of target genes, could determine whether different outputs follow the same ligand equivalence relationships as SMAD1/5/8 phosphorylation (Saxena, Agnihotri, and Sen 2018; Gunnell et al. 2010; Blitz and Cho 2009). Third, a similar approach could be applied to other pathways, such as Wnt or FGF, which also exhibit promiscuous interactions among multiple ligand and receptor variants (Gleason, Szyleyko, and Eisenmann 2006; Kettunen and Thesleff 1998). In the longer term, contextual equivalence groups could function as a ‘periodic table’ that organizes ligands according to their functional effects and contextual interactions in a unified manner to explain why specific components operate together in different developmental contexts and to allow context-specific control of pathway activity for therapeutic applications.

## Supporting information

Key Resources Table

## Acknowledgments

This work was supported by the Defense Advanced Research Projects Agency (contract HR0011-16-0138), the Gordon and Betty Moore Foundation (grant GBMF2809 to the Caltech Programmable Molecular Technology Initiative), the Human Frontiers Science Program (grant RGP0020), the Institute for Collaborative Biotechnologies (grant W911NF-09-0001 from the U.S. Army Research Office), the National Institutes of Health (NIH) (grants R01 HD075335A and R01 MH116508), and the Paul G. Allen Frontiers Group and Prime Awarding Agency (award UWSC10142). This work does not necessarily reflect the position or policy of the U.S. Government, and no official endorsement should be inferred. H.K. is supported by a National Science Foundation graduate research fellowship (grant DGE-1144469). C.J.S. is supported by the NIH National Institute of General Medical Sciences (grant T32 GM008042) and a David Geffen Medical Scholarship. M.A.L. is supported by the National Sciences and Engineering Research Council of Canada Postgraduate Doctoral Scholarship. M.B.E. is a Howard Hughes Medical Institute Investigator.

## Author Contributions

H.K., Y.E.A., and M.B.E. conceived and designed the experiments. H.K., M.A.L., and J.M.L. performed the experiments. H.K. and M.A.L. analyzed the experimental data. H.K., Y.E.A., and C.J.S. developed and fit the model. H.K., Y.E.A., and M.B.E wrote the paper.

## STAR Methods

### Resource Availability

#### Lead Contact

All requests for resources and reagents should be directed to and will be fulfilled by the Lead Contact, Michael Elowitz.

#### Materials Availability

The recombinant proteins, lentiviral particles, siRNAs, and qPCR probes used in this paper are described in Tables S1-S4 and can be purchased from commercial sources, as detailed in the Key Resources Table. Plasmids and cell lines generated for this paper, as indicated in the Key Resources Table, are available from the Lead Contact upon request.

#### Data and Code availability

The datasets and analysis scripts generated used to produce paper figures are available here: https://doi.org/10.22002/D1.1693.

### Experimental Model and Subject Details

#### Tissue culture and cell lines

NMuMG (NAMRU Mouse Mammary Gland cells, female) were acquired from ATCC (CRL-1636). E14 cells (mouse embryonic stem cells, E14Tg2a.4, male) were obtained from Bill Skarnes and Peri Tate. All cells were cultured in a humidity-controlled chamber at 37°C with 5% CO2. NMuMG cells were cultured in DMEM supplemented with 10% FBS (VWR #311K18),1mM sodium pyruvate, 1unit/mL penicillin, 1μg/mL streptomycin, 2mM L-glutamine, and 1X MEM non-essential amino acids. ES cells were plated on tissue culture plates pre-coated with 0.1% gelatin and cultured in standard pluripotency-maintaining conditions (Smith, 2001) using DMEM supplemented with 15% FBS (ES qualified, GIBCO #16141), 1mM sodium pyruvate, 1unit/mL penicillin, 1μg/mL streptomycin, 2mM L-glutamine, 1X MEM non-essential amino acids, 55μM β-mercaptoethanol, and 1000units/mL leukemia inhibitory factor (LIF).

#### Reporter cell line construction

To construct reporter cell lines, a plasmid harboring the BMP response element (Korchynskyi and ten Dijke, 2002) in the enhancer region of a Major Late Promoter (MLP) driving the expression of an H2B-mCitrine protein fusion was integrated into the NMuMG genome using PiggyBac integration (System Biosciences). After transfection, cells were selected with 100μg/mL hygromycin. The base NMuMG reporter was a clonal cell line generated by limiting dilution. This reporter construct exhibited lower background YFP expression and increased signaling dynamic range relative to a similar NMuMG transcriptional reporter (Antebi et al., 2017), consistent with other observations of MLP’s lower background relative to mCMV (Ede et al., 2016). A previously-reported mESC reporter was used (Antebi et al., 2017). To construct this reporter, a related construct, carrying minimal CMV in place of MLP, was randomly integrated into the E14 genome using the FugeneHD reagent and a clonal population selected by colony picking.

### Method Details

#### Quantifying BMP responses with flow cytometry

To assay BMP pathway activity, we genomically integrated a transcriptional reporter into all cell lines as described above (see “Reporter cell line construction”), which expressed YFP proportional to pSMAD activation (Antebi et al., 2017). To measure how a given ligand or combination of ligands activated the pathway, we followed the following protocol in all robotic or manual experiments. Cells were plated at a confluency that, assuming a 24 hour doubling time, would produce sufficient cells for flow cytometry at the experiment endpoint (i.e. at least 5000 cells 36 hours after plating in a 96 well format, or about 20-30% confluent at the start of the experiment). Twelve hours after plating, cell media was removed and replaced with ligand-containing media. Twenty-four hours after ligands were added, cells were lifted from the plate for flow cytometry, by five minute incubation at 37°C in 0.25% Trypsin (or Accutase, for mESCs) following PBS wash. Trypsinization was quenched by resuspending cells in flow buffer, (HBSS containing 2.5mg/mL Bovine Serum Albumin (BSA)). Cells were then filtered through a 40μm mesh and analyzed by flow cytometry (MACSQuant VYB, Miltenyi or CytoFLEX, Beckman Coulter). Note that because of difficulties with cellular aggregation following filtering, alternative flow buffer formulations additionally included either 1mM ethylenediaminetetraacetic acid (EDTA) or 200U/mL DNAse I in some cases.

#### Robotic liquid-handling protocol

For each cell line, a similar protocol was used to generate a full pairwise dataset. First, the transcriptional BMP reporter was generated, and the receptor profile was assayed by qPCR. The only exception was the mESC reporter, whose receptor profile was obtained from a prior publication that assayed receptor expression by RNA-Seq (Antebi et al., 2017). Then, we determined the dynamic range of each ligand in this new receptor context. To do this, we completed at least one biological replicate of single ligand dose responses up to very high concentrations that were expected to approach saturation (e.g. 3μg/mL), with a high fold dilution to capture the full dynamic range. These dose responses fell into three types: saturating activators, non-saturating activators, and non-activators. For saturating activators, we selected a maximum concentration and fold dilution that sampled concentrations corresponding roughly to 25, 50, 75, and 100% of maximal activation. For non-saturating activators, maximal activation was unknown. To avoid supraphysiological concentrations, we limited the maximum concentration to 2 or 3μg/mL and used a minimum 2-fold dilution to sample what little of the dynamic range was accessible. For non-activators, there was no dynamic range. We therefore selected maximum concentrations and fold dilutions that approximated the median dynamic range of the activating ligands, reasoning that these highly homologous ligands had, to a first approximation, reasonably similar concentrations over which they were most active. The full list of all concentrations and fold dilutions used for each cell line is in Table S1.

Next, we assayed all pairwise interactions, with single ligand gradients as well as “pseudo-pairs” of ligands with themselves as controls. For ten ligands, the screen included 10 gradients, 10 “psuedo-pairs,” and the 45 possible pairs, or 65 rows of measurements. Each row included a no BMP control, plus the nine ratios at which each pair (ligand by itself, ligand paired with itself, or ligand paired with another ligand) was assessed. The order of the rows and of the ligands was scrambled between biological repeats.

To expose cells to these complex ligand combinations, we used a custom liquid-handling protocol on the Tecan EVO Freedom 200 base unit, which was housed in a sterile, laminar flow hood. To start the experiment, we manually plated cells eight to twelve hours before the start of the robotic protocol and prepared serial dilutions of the ten ligands by hand. The robotic protocol then added the precise ligand combinations to the appropriate plates, and timed the start of each plate to match the staggered analysis of each plate twenty-four hours after the ligand combinations were added. Specifically, at the appropriate time point, the robot removed a plate from the incubator, aspirated the media, replaced it with the appropriate ligand-containing media, and returned cells to the incubator. The robot incubator was fed 5% CO2. All plates were then collected from the robot incubator at the end of the experiment.

Finally, we quantified cell responses in each of these ligand conditions. 20 to 24 hours after the BMP ligands were added, we measured YFP of each well by flow cytometry, as described above (see “Quantifying BMP reporter with flow cytometry”). We analyzed the resulting flow data with custom MATLAB scripts available at https://doi.org/10.22002/D1.1693, following the steps described below in “Data Processing.”

#### Stable shRNA knockdown and ectopic receptor expression

Individual BMP receptors were knocked down or ectopically expressed in the base NMuMG reporter cell line. For receptor knockdown, lentiviral particles containing constructs for constitutive shRNA expression reported by mCherry (SMARTvector, Dharmacon) were transduced into cells. Per manufacturer’s instructions for determining optimal transduction conditions, we transduced cells at an MOI of 20, using serum-free NMuMG growth media supplemented with 10μg/mL polybrene. 48h after transduction, cells were selected and continuously maintained in 3μg/mL puromycin. Cells with BMPR1A and BMPR2 knockdowns were polyclonal populations generated by a single shRNA. Cells with ACVR1 knockdown were a clonal population selected by limiting dilution of cells transduced with a pool of three shRNAs. For a list of the shRNAs used, see Table S2.

For ectopic expression of BMP receptors, constructs for constitutive expression of mouse receptor cDNAs, reported by mTurquoise, were integrated into the base NMuMG reporter by PiggyBac integration (Systems Bioscience), using previously-reported plasmids (Antebi et al., 2017). Cells were selected and maintained with 500μg/mL geneticin.

#### Transient siRNA knockdown

Cells were plated at 40% confluency in a 24 well plate format with 30μM total siRNA (ThermoFisher Silencer Select #4390771) and 3μL RNAiMAX (Life Technologies). For every gene, a pool of two distinct siRNAs was used, listed in Table S3. Cells were passaged after 24 hr and then used for the relevant experiments.

#### RT-qPCR

Total RNA was harvested from cell lysate using the RNeasy mini kit (QIAGEN), and cDNA was generated from 1μg of RNA using the iScript cDNA synthesis kit (BioRad) per the manufacturer’s instructions. Primers and probes for specific genes (Table S4) were purchased from IDT. Reactions were performed using 1:40 dilution of the cDNA synthesis product with either IQ SYBR Green Supermix or SsoAdvanced Universal probes Supermix (BioRad). Cycling was carried out on a BioRad CFX96 thermocycler using an initial denaturing incubation of 95° for 3 min followed by 39 cycles of (95°C for 15 s, followed by 60°C for 30 s). Each reaction was run at least in duplicate.

### Quantification and Statistical Analysis

#### Analysis of RNA-Seq datasets

To survey BMP ligand and receptor expression across various tissues, we selected relevant datasets from a collection of bulk, mouse RNA-Seq datasets (Lachmann et al., 2018). Datasets that indicated the sample library contained transcriptomic cDNA prepared from polyA RNA and that indicated a tissue-related source were kept. RNA-Seq quantification of BMP receptor expression in the NMuMG parental cell line and the mESC reporter cell line (cf. Figure S3A) was taken from GSE98674 (Antebi et al., 2017).

#### BMP alignment and clustering

To analyze amino acid sequences of BMP ligands, the relevant recombinant sequences (indicated by R&D Biosystems) were globally aligned with MATLAB’s “multialign” function, using default parameters. Distance between sequences was computed by the BLOSUM50 matrix, comparing only positions with no gaps across all aligned ligands, and they were clustered with average-distance linkage.

#### Data processing

All flow cytometry data for gradients and rims was gathered by the same method, implemented manually or robotically (see “Quantifying BMP reporter” and “Robotic liquid-handling protocol” above). The resulting flow data were analyzed in the following manner. Using a custom MATLAB GUI (https://antebilab.github.io/easyflow/), cells were gated on Forward Scatter (FSC) and Side Scatter (SSC), and median YFP used to summarize pathway activation in each well. A custom MATLAB script then dropped any wells with fewer than the threshold number of cells (i.e. 500), reordered wells to match standard ligand order, and subtracted background. For rims, an additional step minimized noise between the eleven plates, which, due to small variation in plate preparation and processing times, had incubation times varying from 20 to 24 hours. Assuming linear accumulation of YFP with time, variation in incubation generates a fold change in background-subtracted YFP. Therefore, each plate was linearly rescaled to a reference plate to minimize total least squares error between technical replicates shared between the two plates. Following standardization of each biological replicate, they were aggregated by another script. To reduce variance due to biological noise, each biological replicate was rescaled by a single linear factor that minimized total least squares error between all corresponding measurements in the two replicates.

To complete gradient analysis, all biological replicates were fit to a Hill function by minimizing total least squares error between the Hill fit and all rescaled biological replicates. To complete rim analysis, Relative Ligand Strength (RLS) and Interaction Coefficients (IC) were computed. RLS is the ratio of each ligand’s maximal activation to the strongest ligand’s maximal activation. However, the IC metric, defined to quantify small differences in pathway activation, required more careful analysis to ensure robustness to technical and biological noise, as described in the following section.

#### Interaction Coefficient

##### Drug synergy classification cannot fully capture ligand interactions

Classification of synergy and antagonism occurs in many fields, including the study of drug interactions and gene epistasis (Costanzo et al., 2016; Meyer et al., 2019; Russ and Kishony, 2018; Yeh et al., 2006; Zimmer et al., 2016). However, these metrics do not have clear analogies for classifying ligand interactions for at least two reasons.

First, all classifications of synergy, when agnostic of mechanism, depend on a phenotypic definition of additivity, i.e. a non-interacting relationship. However, the many definitions of additivity developed for drug interactions are not consistent with each other and not necessarily appropriate to describe ligands activating the same pathway. As an example of this, most definitions of additivity descend from approaches defined by Loewe (Loewe, 1928) and Bliss (Bliss, 1939), who predicted that additive outcomes would be the sum of each component’s dose or effect, respectively (Russ and Kishony, 2018). However, these assumptions make little sense in the context of two signaling proteins that use the same signaling components to produce their individual effects; indeed, individually saturating concentrations are not expected to sum in their effects or their normalized dose. A third approach to additivity considers the effect of each drug as an independent event, such that a non-additive response deviates from the known joint probability of independent outcomes (Yeh et al., 2006). But the sharing of pathway components between the two ligands makes this assumption of independence inappropriate.

Second, classifications of interaction type depend on well-defined regimes of behavior. The effect of drug combinations on growth rate has distinct boundaries for these regimes, but there are not clear boundaries for pathway activation. For example, the largest effect of a drug is to decrease growth rate to zero, but the upper bound of a signaling protein’s effect could be set by any one of the pathway components being limiting and so depends heavily on cell context. Similarly, drugs can exhibit a “strong” form of antagonism when a combination causes cells to grow faster than in the absence of drugs. By contrast, signaling proteins cannot, in combination, decrease pathway activation below zero, i.e. the response in the absence of signaling proteins. Thus pathway activation requires a different separation between weak and strong antagonism.

##### Interaction Coefficient captures features specific to ligand interactions

To address the incongruency of drug and ligand interactions, we developed the Interaction Coefficient, a definition of additivity and associated regimes of behavior that are appropriate for ligand interactions as described above. The goal of defining this metric was to quantify and compare specific ligand behaviors of interest, rather than providing a generic or alternative synergy metric to an increasingly large list (Russ and Kishony, 2018; Tekin et al., 2016; Vlot et al., 2019). IC focuses on a definition of additivity most relevant for this study: additively interacting ligands combine with each other in the same way they combine with themselves.

At first, this presents an apparent challenge, as the way a ligand combines with itself is inconsistent. In the limit of very low and very high concentrations, doubling ligand concentration has no effect on pathway output. Conversely, at intermediate concentrations, doubling ligand concentration can increase pathway output by two-fold or more, depending on the cooperativity of the response. Rather than making the strong claim that any one of these behaviors is the true additive expectation, IC includes both as boundaries between distinct behavioral regimes, where output greater than the sum is synergy and output less than the maximum of the two is antagonism of some strength. Finally, rather than predicting which of these two behaviors would result when a ligand was added with itself, we simply measured them. Thus, while IC_AB_ describes the phenotypic response when ligand *A* combines with ligand *B*, ΔIC, or the difference between IC_AA_ and IC_AB_, indicates whether any apparent non-additivity between *A* and *B* is truly unexpected.

To complete our definition of IC, we add a third boundary dividing weak and strong antagonism, determined by whether the combination response interpolates or is less than both individual responses. Thus, the three boundaries define four behavior regimes. Within each regime, the strength of that behavior type can be computed by normalizing the observed response to the maximal effect size. This generates the following definition of IC:

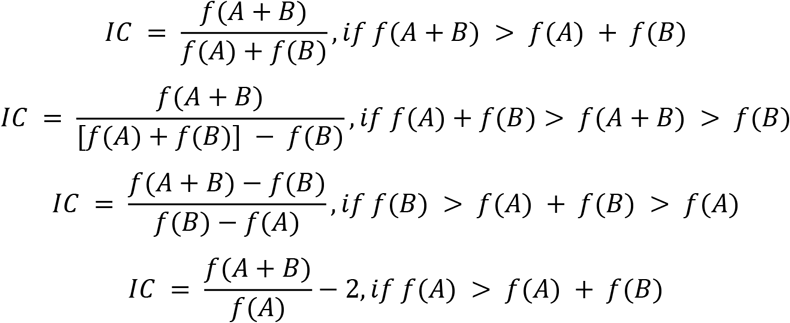

where *f(X)* is the nonnegative pathway response to ligand input X, and *f(B) > f(A)*. Note that IC has no upper bound, consistent with the lack of upper bound on pathway activation, and that this calculation assumes all data have been background subtracted.

However, this definition of IC misclassifies small absolute changes in *f(A+B)* as strong pairwise interactions in a few cases. Specifically, if *f(A)* ≈ 0 or *f(B)* ≈ *f(A)*, the denominator of IC diverges, respectively, in the additive and antagonistic regimes, effectively due to the collapse of one of the behavior regimes.

We therefore defined a discontinuous definition of IC, which normalizes each regime to a more intuitive standard that does not approach zero, which is the response to the stronger ligand individually, or *f(B)*. The Interaction Coefficient (IC) as reported in this paper is computed with the following piecewise formula:

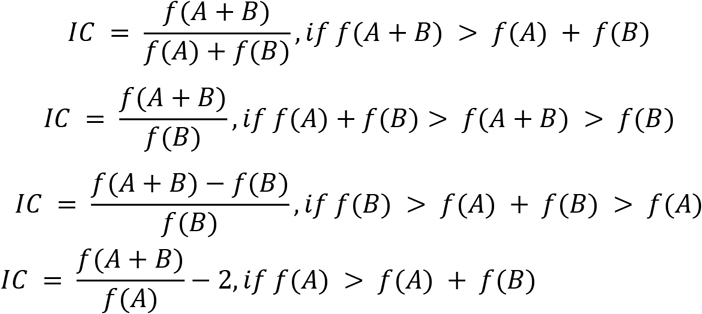

where *f(X)* is the nonnegative pathway response to ligand input X, and *f(B) > f(A)*, as above. To describe the interaction of any ligand pair, the largest magnitude IC value, positive or negative, for all ratios of A to B is reported.

##### Determining the Interaction Coefficient from noisy data

Because *f(X)* is sampled only three or four times from a broad, underlying distribution, we sought a robust metric to determine if two sets of measurements, *f(X)* and *f(Y)*, could be reasonably considered different from each other, in both a statistical and biological sense. Our first criterion was that the range of replicates of *f(X)* must not overlap with the range of *f(Y)* replicates. However, even technical replicates collected on different days (i.e. *f(X)* and *f(X’)*) were sometimes non-overlapping, indicating these measurements were distributed over a large range. Therefore, we added an additional criterion that the degree of non-overlap must exceed a certain threshold. Many of the apparently spurious classifications occurred for low YFP values that were a large percentage change in background but an infinitesimal percentage of the overall dynamic range. Therefore, we set a threshold for effect size, which was the maximum, background-subtracted YFP observed in a given cell line’s full dataset divided by 30, i.e. ~3% of the cell line’s dynamic range. This threshold achieved two objectives, nearly eliminating spurious identification of strong pairwise interactions due to small variations in background and most closely matching a manual classification of pairwise interaction type.

Nonetheless, the correct, automatic classification of some rims was not possible with this approach, perhaps due to a liquid-handling error specific to one or a set of measurements. For these 13 of the 385 total pairs (<4% of the full dataset of 7 cell lines with 45 pairs and 10 controls pairs each), we determined a more reliable estimate of IC value by selecting a less noisy point on the rim, calculating it from independent measurements of that same pair, or setting it to 0 for the null hypothesis of additivity. A full description of these pairs and their manual classification is included below.

##### Rim classification corrections

Confident, automatic classification of all pairwise interactions posed a significant challenge. Measurements were limited to three or four biological replicates while small robotic errors, such as dropped tips or cross-well contamination, randomly disturbed a small subset of the data. While the analysis pipeline outlined above minimized effects of technical and biological noise by rescaling, and randomization of experiment layout distributed any systematic errors more evenly, the automatic classification of pairwise interactions did not match our intuition in a small number of cases (13 of the 385 rims). Here we describe those cases, why they resisted automatic classification, and how we manually corrected the classification.

First, for two measures of pathway activation to be considered different, the ranges of their respective biological replicates had to be non-overlapping. However, a single outlier among three or four points may cause these ranges to overlap, despite significant differences in the median. Thus, plausible non-additive interactions may be classified as additive by one erroneous measurement. For example, in NMuMG cells, BMP2 and BMP4 combined with GDF5 or GDF7 produced antagonism distinct from GDF6’s lack of effect of BMP2 (Figure S2D). Similarly, in mESC, BMP2 and BMP4 interacted very differently with BMP10 than with other activating ligands, and BMP2 was antagonized by GDF5, but not GDF6 (Figure S3E). However, despite these qualitative differences, all these rims were classified as additive due to a single outlier. For correct classification of these rims, the requirement for non-overlap was removed for the specific points on the rim where the pairwise interaction was clear (cf. asterisks in Figure S2D).

Second, a few rims were classified as strong synergy due to a small increase in pathway activation at subsaturating concentrations, though synergy manifesting exclusively at extreme ligand ratios or small absolute effect sizes, both of which were unlikely for true synergistic interactions. Moreover, given the width of the distribution of median YFP measurements, multiple samples from the same distribution could often, by random chance, produce non-overlapping measurements. For example, In NMuMG cells, despite the similarity of responses to BMP9 paired with GDF5 or GDF6, the latter was classified as synergy due to lack of overlap at low activation levels (Figure S2E). Similarly, in the BMPR2 knockdown cell line, BMP2 paired with BMP5, BMP6, and BMP7 showed synergy at low activation levels, despite not much clear difference from BMP4’s additive interaction with BMP5 (Figure S5G). These small variations were not considered sufficient evidence for a strong claim of synergy, and these pairwise interactions were classified as additive, the null hypothesis for all pairs.

Finally, two unexpected edge cases emerged in the ACVR1 knockdown. The combination of BMP4 with BMP9 generated both positive and negative IC values, though the positive IC values were larger (Figure S4D). However, the negative IC effect was larger in absolute terms and consistent strong antagonism observed in similar rims, such as BMP5 with BMP9. Therefore, this pair was summarized by only its largest negative IC value. Lastly, BMP2 with BMP10 appeared to be a borderline suppressive interaction. A closer examination of this pair at higher concentrations and more ligand ratios confirmed the suppression between BMP2 and BMP10, and this IC value was used to classify these ligands (Figures S4E,F).

##### Interaction Coefficient provides useful classification and agrees with existing methods

The Interaction Coefficient (IC) enabled several parts of the analysis presented here. First, as a dimensionless quantity, it allowed easy comparison between cell lines with drastically different dynamic ranges, either due to differences in sensor behavior or data being collected on different cytometers. Second, as a quantitative, rather than qualitative, metric, IC allowed “softer” classification of pairwise interactions, since small amounts of noise could possibly generate an observation of a weak, but not strong, antagonistic interaction. Third, IC reliably distinguishes distinct behavior types for a wide array of individual ligand doses, as summarized in Figure S1B. The few difficulties of using the IC do not limit its usefulness in this study. One possible drawback to computing IC as described here is the discontinuity of the metric, introduced to avoid divergence when *f(A)* = 0 or *f(A)* ≈ *f(B)*. However, these discontinuities preserve better our intuitive classifications of antagonism and suppression. Another challenge is that IC is computed by sampling many ligand ratios, computing IC for each ratio, and then summarizing the pairwise behavior with the largest magnitude IC value, whether positive or negative. However, this is a feature of all synergy metrics, which are defined with respect to the concentrations and ratio at which the pair is studied. Rather, this highlights the importance of reporting synergy alongside the concentration regime that was sampled and recognizing that all estimates of non-additivity represent lower bounds.

Ultimately, IC agrees with preexisting synergy metrics, while providing improvements specific to this application. Defining a synergy metric as distinct regimes normalized to their maximum effect size as implemented by Yeh *et al*. (2006), provides the basis for IC, though with a unique definition of additivity. Similarly, IC summarizes the same features described by Relative Ligand Strength and Ligand Interference Coefficient in another study of BMP pairwise interactions (Antebi et al., 2017), but with two small differences. First, an RLS value less than one previously classified an interaction as ratiometric (i.e. antagonistic), but our study showed that not all non-activating ligands (e.g. RLS ~ 0) antagonized activating ligands, as captured by IC’s comparison of the combination response to activation by the stronger ligand alone. Second, nonzero LIC values earn a classification of imbalance or balance, but depend on the assumption that each ligand is individually saturating, such that any increase or decrease relative to that amount is noteworthy. However, as observed in mESCs (Figure 4A), not all ligands appeared to saturate the pathway. The assumptions underlying IC allow its use at subsaturating ligand concentrations, as in the case of BMP9 in mESCs.

#### Determining equivalence groups

To classify the functional similarity of ligands, we sought a quantitative method to summarize differences between each ligand’s pattern of individual and pairwise activation. To this end, we used hierarchical clustering to summarize the hierarchy of possible ligand groupings. First, we computed ligand differences as the Euclidean distance between each ligand’s eleven features in a given cell line (i.e. nine IC values in pairs with the nine other ligands, one IC value in pair with itself, and one RLS value). RLS is multiplied by two for reasons described below. Next, we agglomeratively clustered ligands with complete linkage, such that the differences between two clusters is equal to the distance between their two most dissimilar members. Finally, we cut the resulting dendrogram to generate “monochromatic clusters” (cf. (Segrè et al., 2005)), such that ligands in the same group had similar activation strengths and had the same interaction type (i.e. color of synergistic, additive, antagonistic, suppressive), though not necessarily interaction strength, with all other ligands. The definition of our metrics allowed us to define a distance threshold of 1, which roughly corresponds with the desire for monochromaticity. Specifically, a distance of 1 corresponds with a difference of 1 in IC, which is the distance between the different regimes of IC behavior, or corresponds to a change of 0.5 in RLS (when RLS has been multiplied by 2), which is the difference between a strong and weak activator. Thus, most groups are defined by clusters at a distance of 1, with some small adjustments to preserve monochromaticity.

##### Global equivalence groups

When determining global equivalence groups, each ligand has seven times as many features (i.e. IC and RLS values) than when determining equivalence in a single cell line. To maintain a similar threshold, we increased the squared distance threshold seven-fold from 1 to 7, and cut the tree at a Euclidean distance of 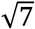. The five resulting equivalence groups are in fact more robust to choice of distance than other possible groupings, as they are consistent with the widest possible range of thresholds.

This threshold increase allows ligands in the same global equivalence group to have small differences in pairwise interactions, meaning the global equivalence groups are not strictly “monochromatic.” However, since a claim of global inequivalence is a stronger claim than inequivalence in a single cell line, this reflects our intuition that a larger pattern of differences is required to assign ligands to distinct groups and reduces the influence of a single pairwise measurement on the global classification. This is important because the three ligands whose global equivalence would be most affected by lowering the threshold, GDF5, GDF6, and GDF7, had particularly challenging to identify pairwise interactions, as a result of being non-activators.

Specifically, GDFs were placed in separate equivalence groups only when they differed in their ability to produce antagonistic interactions (NMuMG cells, mESCs, ACVR1 knockdown, ACVRL1 ectopic expression) or when they differed in strength of antagonism (BMPR1B ectopic expression). However, a ligand’s inability to produce an antagonistic interaction may not reflect its intrinsic properties, but instead indicate that it is present at too low a concentration. For non-activating ligands, it is difficult to determine which concentrations are sufficiently high and, in particular, whether all non-activators should be used at the same concentration. Indeed, GDF dose-responses in the single cell context where they activated, BMPR1B ectopic expression, showed significant differences in the three ligands’ EC_50_ values (Figure S6E, inset). Therefore, before confidently placing GDFs in separate global equivalence groups, the lack of antagonism or suppression in some cases should be confirmed by repeating experiments with more diverse GDF concentrations.

#### BMP Mathematical Model

##### Model assumptions and justification

Dimeric BMP ligands activate the BMP pathway by complexing with two Type I and two Type II receptors. Once the complex is assembled, constitutively active Type II receptors phosphorylate and activate Type I receptors, which enzymatically activate SMAD1/5/8 molecules, generating a pool of pSMAD that translocates to the nucleus to promote gene expression. To focus on a minimal number of pathway components and steps, our model of BMP pathway activation considered only one-step formation of a trimeric signaling complex, containing a ligand, a Type I receptor, and a Type II receptor. In the model, each ligand binds each receptor dimer with a unique affinity and activates with a unique activity. Ligand specific-activity is a well-documented phenomenon in BMP signaling, though the mechanism is not clear (Nickel and Mueller, 2019).

This simplified model neglects certain details of BMP pathway activation, which decreases model complexity without significant loss of explanatory power for our *in vitro* system. Specifically, one-step complex formation overlooks the stepwise assembly of BMP signaling complexes. However, summarizing a two-step binding process with a single phenomenological parameter does not limit the possible pairwise interactions allowed by the pathway architecture (Su et al. 2020). The reduced model also neglects the pentameric nature of the full BMP signaling complex, where one BMP dimer binds two Type I and two Type II receptors. This model also does not include the other secreted and membrane-bound molecules that can influence signaling complex formation, such as secreted ligand antagonists (e.g. Twsg1, Noggin, Chordin), co-receptors (e.g. RGMA, ENG), pseudo-receptors (e.g. BAMBI), or other inhibitors (e.g. Fst). However, siRNA targeting the most highly-expressed of these molecules in NMuMG, including Twsg1, Fst, and Rgmb, had no effect on pairwise interactions in NMuMG (Antebi et al., 2017).

##### Model equations and solution

To model BMP pathway activation, we used mass-action kinetics to describe the one-step formation of tripartite signaling complexes (*T_ijk_*), where one ligand (*L_i_*) binds one Type I receptor (*A_j_*) and one Type II receptor (*B_k_*) with a complex-specific affinity (*K_ijk_*). Each complex also has a distinct activity (*ε_ijk_*) with which it phosphorylates downstream SMAD proteins, and each complex contributes to the overall measured output (*S*). We assume that binding kinetics reach steady-state more rapidly than complexes activate the pathway. We also assume that ligand concentration, provided by a large reservoir of cell media in experiments, is effectively constant. As shown in Su *et al*. 2020, this results in the final equations:

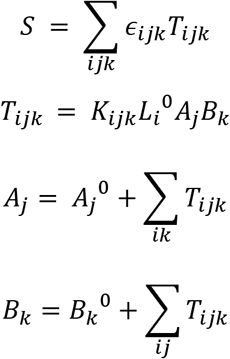

where 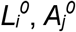 and 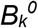 correspond to the starting concentrations of these free species. Thus, when the receptor and ligand environment are specified (i.e. 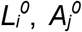, and 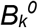 are known), pathway activation can be solved for, as a function of independent variables are *K_ijk_* and *ε_ijk_*. The implicit dependence of *S* on the model parameters can be solved by constrained optimization, implemented by the Python package *promisys* (Su et al. 2020).

##### Parameter fitting

To maximally constrain the fit parameters, we selected datasets that could be fit by the fewest parameters, which scale with the number of ligand and receptor components. To this end, the model was fit to five ligands signaling in cell lines that could be modeled with only five of the seven BMP receptors. Specifically, we focused on BMP4, BMP7, BMP9, BMP10, and GDF5, representing each one of the global equivalence groups and neglecting possible differences between GDFs. We further selected the NMuMG and receptor knockdown datasets, which express the fluorescent BMP reporter from the same genomic locus and express at most five BMP receptors. Receptor expression in these four cell lines was also measured by the same RT-qPCR protocol.

As indicated in the model, our measurements of BMP signaling can thus be modeled by *S* and its dependence on 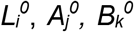, the known ligand and receptor environments, and *K_ijk_* and *ε_ijk_*, the unknown affinities and activities that are fit. However, the data carry various units that also appear in the fit parameters. *S* is measured in units of background-subtracted median YFP, 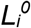 in units of ng/mL, and 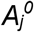 and 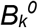 in relative expression units from RT-qPCR abundance normalized to a housekeeping gene. The fit parameters units depend on these values in the following way:

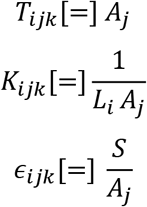

The implicit units in the fit parameters influenced the parameter fitting in the following two ways. First, any global transform of *S*, 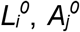, or 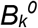 affects absolute parameter values to reflect the change in units, but does not affect the overall fit. Therefore, for convenience, all values of *S*, 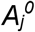, or 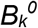 were normalized to a maximum of 1, dividing by the maximum YFP or relative expression value, respectively, across all four cell lines. Second, the lack of meaningful units for the *K_ijk_* and *ε_ijk_* parameters made it difficult to define biologically-reasonable constraints on the absolute values of these parameters. In principle, the values of *K_ijk_* and *ε_ijk_* may take any nonnegative value, but they cannot physically vary over an arbitrary number of orders of magnitude. Therefore, *K_ijk_* and *ε_ijk_* were bounded within six orders of magnitude. To find reasonable upper and lower bounds, a subset of the data was fit with strictly positive *K_ijk_* and *ε_ijk_* values, which used a method similar to the one described below but did not constrain parameters and produced dozens of equivalently good solutions. These sets of solutions showed most best fit parameter values for *K_ijk_* and *ε_ijk_* ranged between 10^−4^ and 10^2^ when *S* and receptor expression were normalized between 0 and 1.

A final consideration in model fitting was the nature of RT-qPCR measurements, where different gene targets are amplified with different probe sets with different efficiencies. These values introduce significantly larger error than 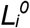, which is directly controlled, and *S*, which is a median across hundreds of cells. At the high C_t_ values of these relatively low abundance receptors, estimates of the same level of expression could vary as much as 10-fold, assuming a 10% difference of efficiency amplified over 25 cycles. As relative receptor levels are crucial in the model, a rescaling parameter, bounded by a 10-fold increase or decrease, was added for each receptor, to reflect the uncertainty of the RT-qPCR estimate of gene expression. Thus, the values passed to the model are instead:

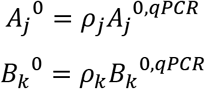

where *A_j_*^0,*qPCR*^ and *B_k_*^0,*qPCR*^ are total receptor expression measured by RT-qPCR and *ρ_j_* and *ρ_k_* are their respective rescaling parameters. To fit *K_ijk_* and *ε_ijk_*, we used SciPy’s least squares function to minimize the error between observed and model-predicted *S*, when 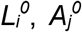, and 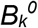 were specified and *K_ijk_, ε_ijk_, ρ_j_*, and *ρ_k_* freely varied over the ranges described above. Each parameter fitting run was initiated with a distinct, randomly-chosen initial condition and resulted in an equal number of distinct solutions, each with some total error. Filtering out solutions whose error changed the type, rather than strength, of pairwise interactions between ligands, we identified a set of 22 solutions that most closely matched the data’s qualitative behavior (Figure S8B). The values for the fit parameters shown in Figure S8D-J are reported in Table S5.

## Supplemental Information Titles and Legends

**Figure S1:**
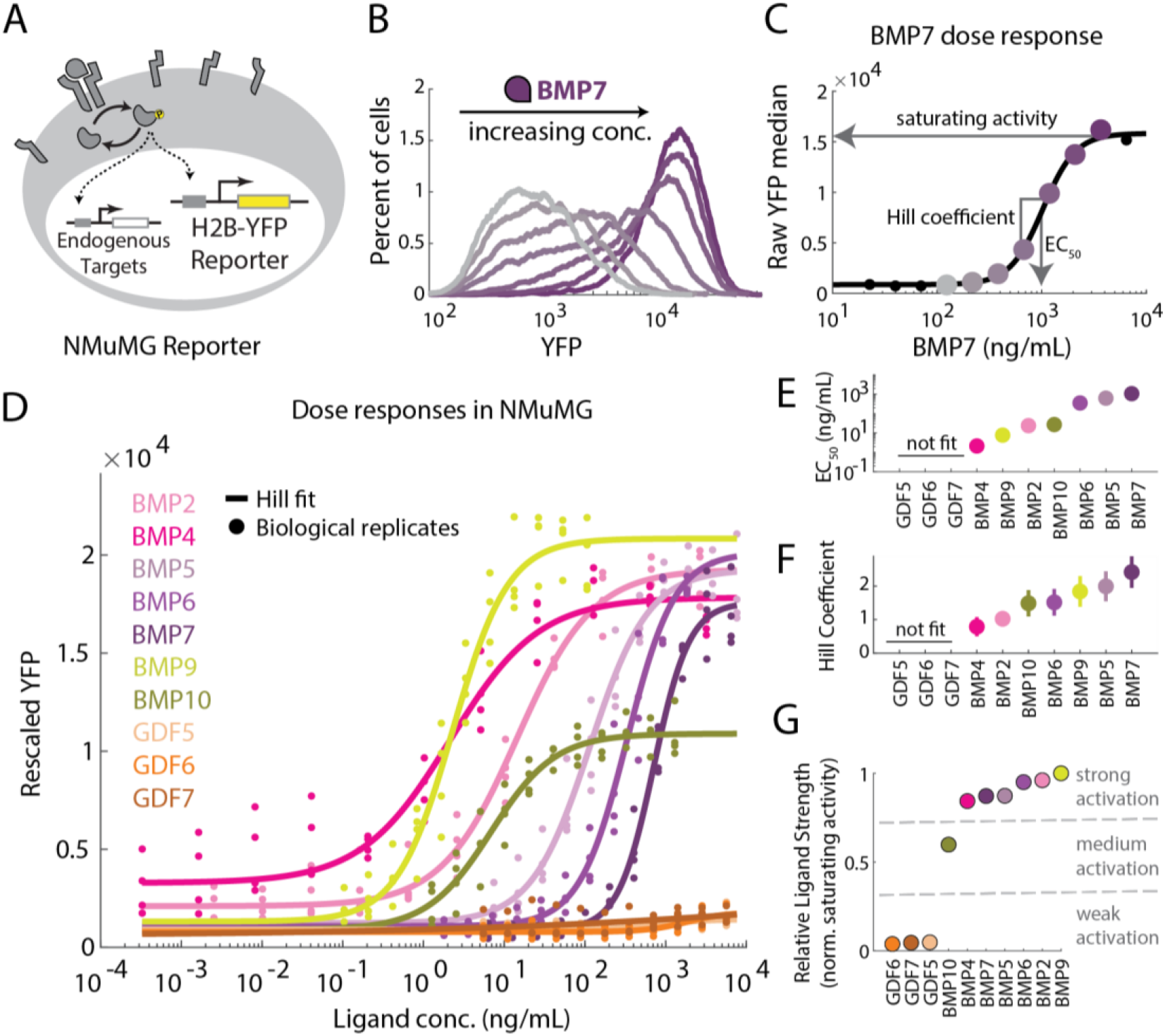
Quantitative dose-responses reveal individual ligand differences. (A) NMuMG reporter cells include a stably-integrated BMP response element driving expression of Histone 2B-mCitrine (H2B-YFP). (B) Flow cytometry data (YFP channel) for a BMP7 dose response in the NMuMG reporter cell line shows concentration-dependent increase in YFP. Colors shaded from gray to purple indicate increasing concentrations of BMP7, plotted in C. (C) Medians of YFP distributions in B show BMP7 dose response, plotted as a function of BMP7 concentrations, which are colored by the same scheme as in B. Black line shows Hill fit of this single biological replicate. Black dots are responses at concentrations not shown in panel B, but acquired in the same manner. (D) Dose responses for ten BMP ligands in the reporter line vary in slope, EC_50_, and saturating activity. Dots represent three biological repeats, and lines indicate the best fits to Hill functions. (E) EC_50_ values from best fit Hill functions for all activating ligands vary continuously over two orders of magnitude. Parameters for GDF5, GDF6, GDF7, whose weak responses do not resemble Hill fits, were omitted. (F) Hill coefficients for activating ligands vary over a small range. (G) Relative Ligand Strength (RLS, saturating ligand activity normalized to the strongest ligand’s saturating activity) values show into weak, medium, and strong activation levels.

**Figure S2:**
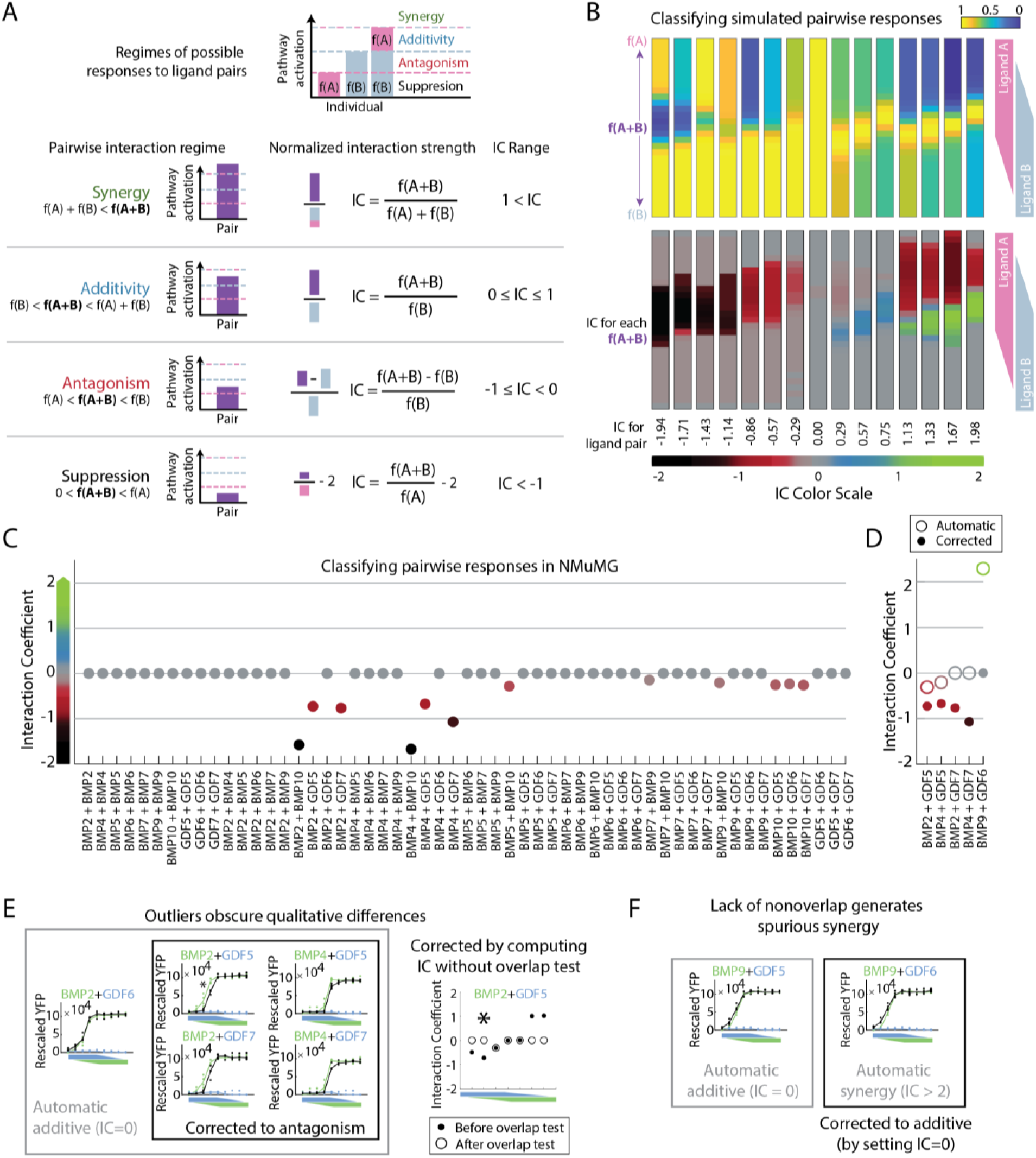
Interaction Coefficient (IC) classifies the range of pairwise interactions observed in simulations and experiments. (A) The Interaction Coefficient (IC) is computed by a discontinuous, piecewise formula corresponding to four regimes of possible pairwise response. These four regimes (synergy, additivity, antagonism, suppression) are defined by the magnitude of pairwise response relative to the individual responses (i.e. f(A+B) relative to f(A) and f(B)). (B) Variety of theoretical measurements produced by a mathematical model of BMP signaling can be ordered and classified by IC. Top row shows theoretical pathway activation by ligand ratios ranging from pure Ligand A to pure Ligand B. Colors indicate low (blue) and high (yellow) values. Bottom row shows corresponding IC values for each theoretical observation, colored by the IC color scale. Theoretical rims are ordered from largest negative interaction to largest positive interaction. (C) For each ligand pair, one IC value is computed (STAR Methods). Most responses are saturated additive (gray), with limited instances of antagonism (red) with GDF5 and GDF7 and suppression (black) with BMP10. (D) Some rims required a different calculation of IC. The automated (open circle) and corrected (closed circle) values are shown for the five pairs where these changes were made. See STAR Methods for full discussion. (E) To be classified as antagonistic, the full set of biological repeats for a rim response must not overlap with the distribution of measurements for the stronger individual ligand. However, for four of the corrected rims, single outliers created overlap and obscured clear differences between the rim and gradient. For example, BMP2 combined with GDF6 clearly overlaps with the response to BMP2 alone and is additive, while BMP2 and BMP4 combined with GDF5 and GDF7 appear qualitatively different. Compare inset plots of pairwise responses, each of which includes two single ligand responses (plotted in blue and green) and the response to the pair (plotted in black). The x-axis corresponds to multiple ratios of the two ligands, as in A. To correctly classify these rims, the requirement for nonoverlap was neglected. As an example, the IC value for each combination of BMP2 with GDF5 is shown, where the ratio with the strongest antagonism (marked by an asterisk) can be recovered by ignoring the overlap test. (F) Similarly, nonoverlap between the spread of points for rim and gradient measurements spuriously classifies rims as synergistic, even though very similar looking rims show significant overlap. BMP9 combines with GDF5 and GDF6 in qualitatively similar ways, despite small differences in whether every point on the rim overlaps with the BMP9 gradient. Thus, BMP9 with GDF6 was classified as additive, to match BMP9 with GDF5. See also Figure 3.

**Figure S3:**
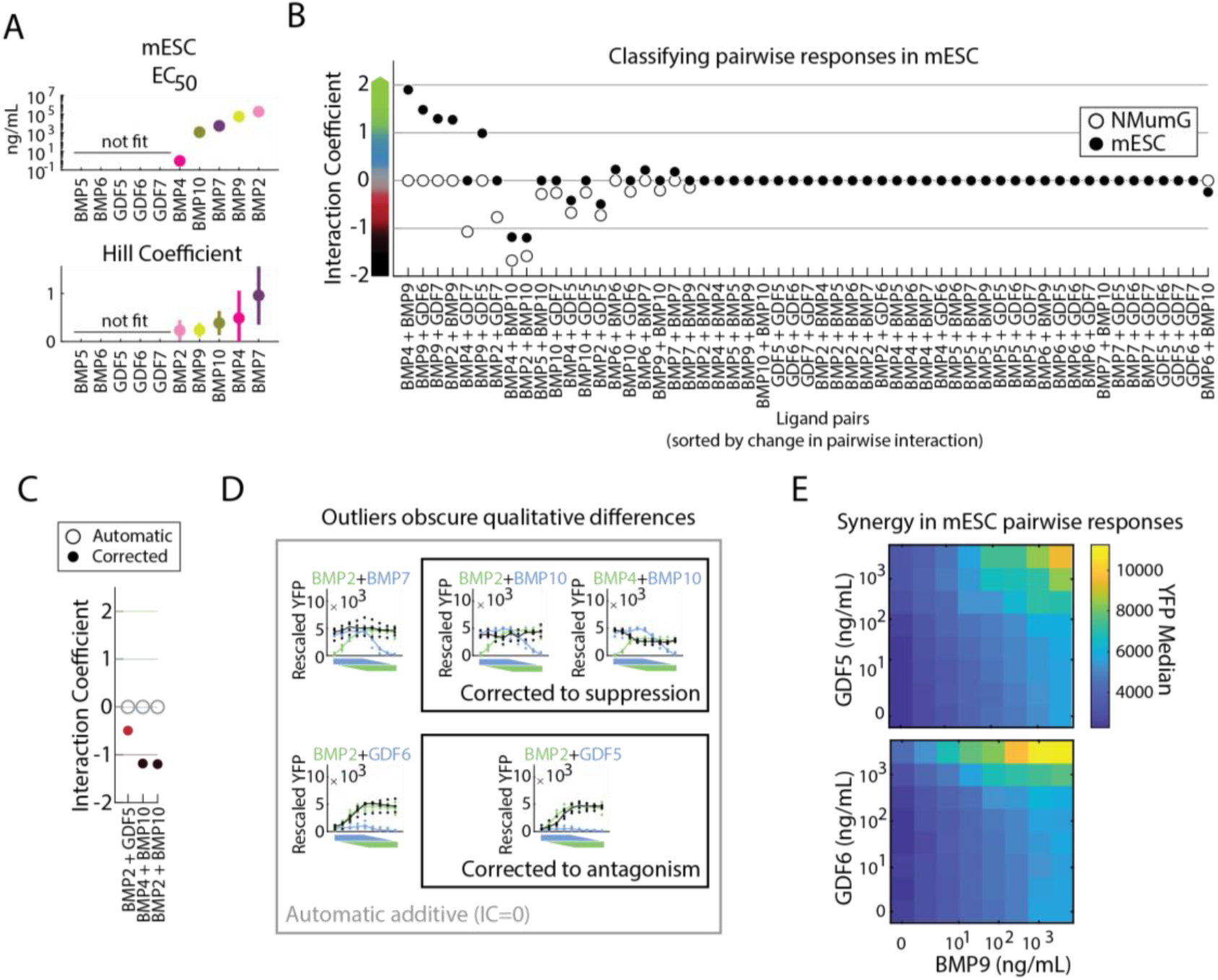
Full quantification of BMP signaling in mESC reveals unique ligand properties. (A) Dose responses to individual BMPs in mESCs were fit to Hill functions to determine best fit parameters and compare ligand behavior. Ligands that do not activate (GDF5, GDF6, GDF7) or do not saturate (BMP5, BMP6) were not fit. Relative to NMuMG cells, Hill coefficients are much lower for mESCs, and EC_50_ values are much higher. (B) Between NMuMG cells and mESCs, a small percentage of pairwise interactions changed significantly, and most changes represented an increase in IC. IC values for all ligand pairs activating mESC (closed circle) and NMuMG cells (open circle, as in S2B) are sorted by decreasing change in IC values. (C) Automatic classification of pairwise interactions failed for three rims, where antagonistic and suppressive interactions were misclassified as additive. See STAR Methods for full discussion. (D) As in Figure S2D, single outliers created overlap between rims and gradients. The corrected IC was computed by ignoring the overlap test for these rims. To show differences between correctly and incorrectly classified pairs, additive interactions of BMP2 with BMP7 or GDF6 are contrasted to BMP2 with BMP10 and GDF5, as well as BMP4 with BMP10. (E) High-resolution matrices of BMP9 with GDF5 and GDF6 in mESC confirm synergy of BMP9 with non-activating ligands. See also Figure 4.

**Figure S4:**
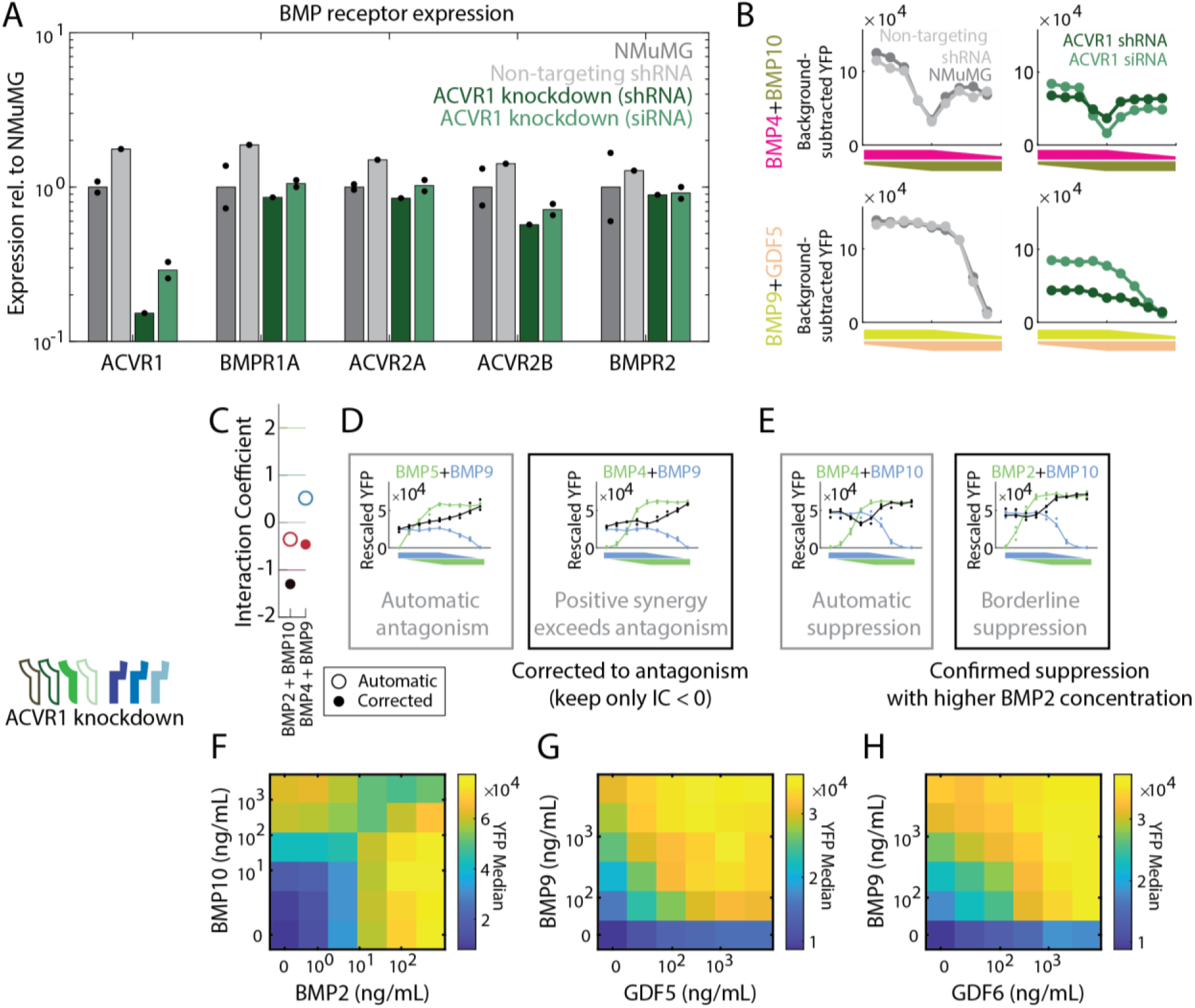
Knockdown of ACVR1 by shRNA phenocopies siRNA knockdown and alters pairwise interactions. (A) RT-qPCR of BMP receptor expression shows potency and specificity of shRNA knockdown of ACVR1 in parental NMuMG cells. ACVR1-targeting shRNA, or transiently transfected with ACVR1-targeting siRNA. shRNA knocks down ACVR1 mRNA more potently than and as specifically as siRNA, while non-targeting shRNA appears to affect all receptors equally. Expression values are normalized to housekeeping gene expression, and then to the average expression of the same gene in NMuMG cells. Dots are biological replicates, and bar height is either one biological replicate or geometric mean of two replicates. (B) Responses to ligand pairs sampled at multiple ratios and overall high concentration for same cell types as in A show that non-targeting shRNA has no effect on pairwise responses, whereas knocking down ACVR1 by shRNA or siRNA produces similar effects, with lower activation by BMP4 and BMP9. (C) Automatic classification of pairwise interactions failed for two rims, as shown in D and E. See STAR Methods for full discussion. (D) BMP4 combined with BMP9 produced a response with synergistic and antagonistic qualities. However, the synergy is a small effect relative to the antagonism, and the rim more closely resembles another strong antagonistic interaction, BMP5 with BMP9. The corrected classification therefore keeps the IC values corresponding to antagonism. (E) BMP2 with BMP10 appeared nearly suppressive, mirroring BMP4 with BMP10. Measuring pairwise interactions at a higher BMP2 concentration revealed a strong suppressive interaction (F), where heatmap intensity indicates pathway activation, reported by YFP median. Similarly, sampling responses to BMP9 with GDF5 (G) or with GDF6 (H) confirmed the balance interactions between these ligands. X- and y-axes in F-H correspond to logarithmically increasing ligand concentration, and colors show pathway activation as reported by YFP median. See also Figure 5.

**Figure S5:**
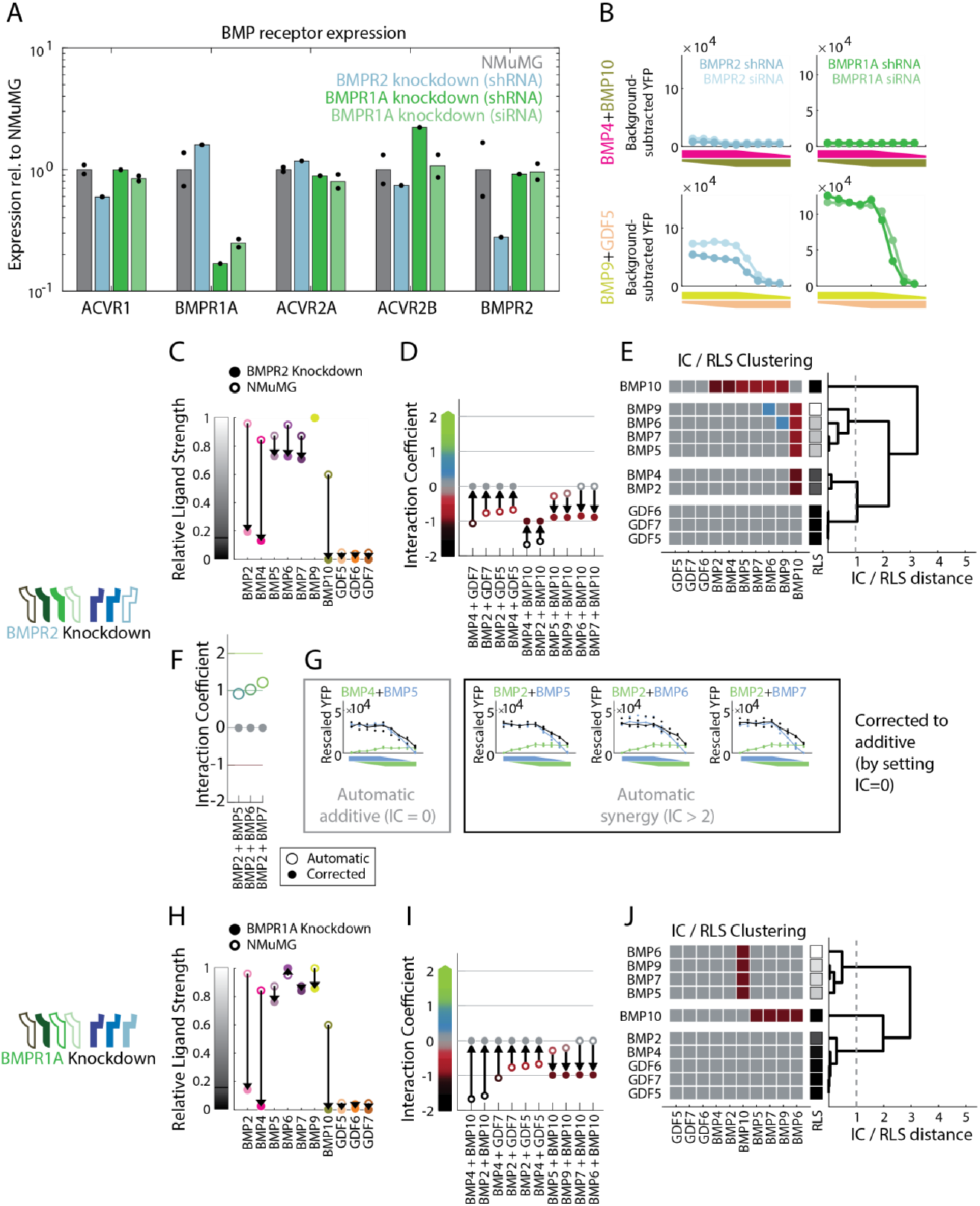
Constitutive shRNA expression knocks down BMPR2 and BMPR1A to produce receptor-specific effects distinct from ACVR1 knockdown. (A) RT-qPCR of BMP receptor expression shows that constitutively expressed shRNA knocks down BMPR2 and BMPR1A potently and relatively specifically, though with small possible increases in BMPR1A and ACVR2B, respectively. Expression values are normalized to housekeeping gene expression, and then to the average expression of the same gene in NMuMG cells. Dots are biological replicates, and bar height is either one biological replicate or geometric mean of two replicates. (B) Despite apparent perturbations to off-target BMP receptors, shRNA knockdown produces qualitatively similar responses to ligand pairs as the more specific siRNA knockdown. Pairwise responses are sampled at multiple ratios and overall high concentration in NMuMG reporter with BMPR2 and BMPR1A targeted by siRNA and shRNA. RLS and IC values change in specific ways following BMPR2 knockdown (closed circle), compared to NMuMG cells (open circle). (C) Activation by BMP2 and BMP4 decreases significantly, while BMP10 activation is lost. (D) Antagonism of BMP2 and BMP4 with GDFs is lost, while all BMP10 interactions become strong antagonism. Changes of IC less than 0.5 are not shown. (E) Hierarchical clustering of all IC and RLS values reveals equivalence groups following BMPR2 knockdown. (F) Automatic classification of pairwise responses failed for three ligand pairs in the BMPR2 knockdown dataset, as shown by the automatic (open circle) and corrected (closed circle) measurements. (G) Small nonoverlap between the rim and the gradient is classified as a synergistic interaction, analogous to Figure S2E. The IC values for BMP2 combined BMP5, BMP6, and BMP7 are corrected to 0, to match the qualitatively similar pair, BMP4 with BMP5. RLS and IC also change following BMPR1A knockdown (closed circle) relative to NMuMG cells (open circle). (H) Activation by BMP2 is decreased while activation by BMP4 or BMP10 is lost. (I) All interactions of BMP2 and BMP4 with other non-activating ligands become additive, while BMP10 strongly antagonizes all activating ligands. Changes of IC less than 0.5 are not shown. (J) Hierarchical clustering of all IC and RLS values reveals equivalence groups following BMPR1A knockdown. See also Figure 6.

**Figure S6:**
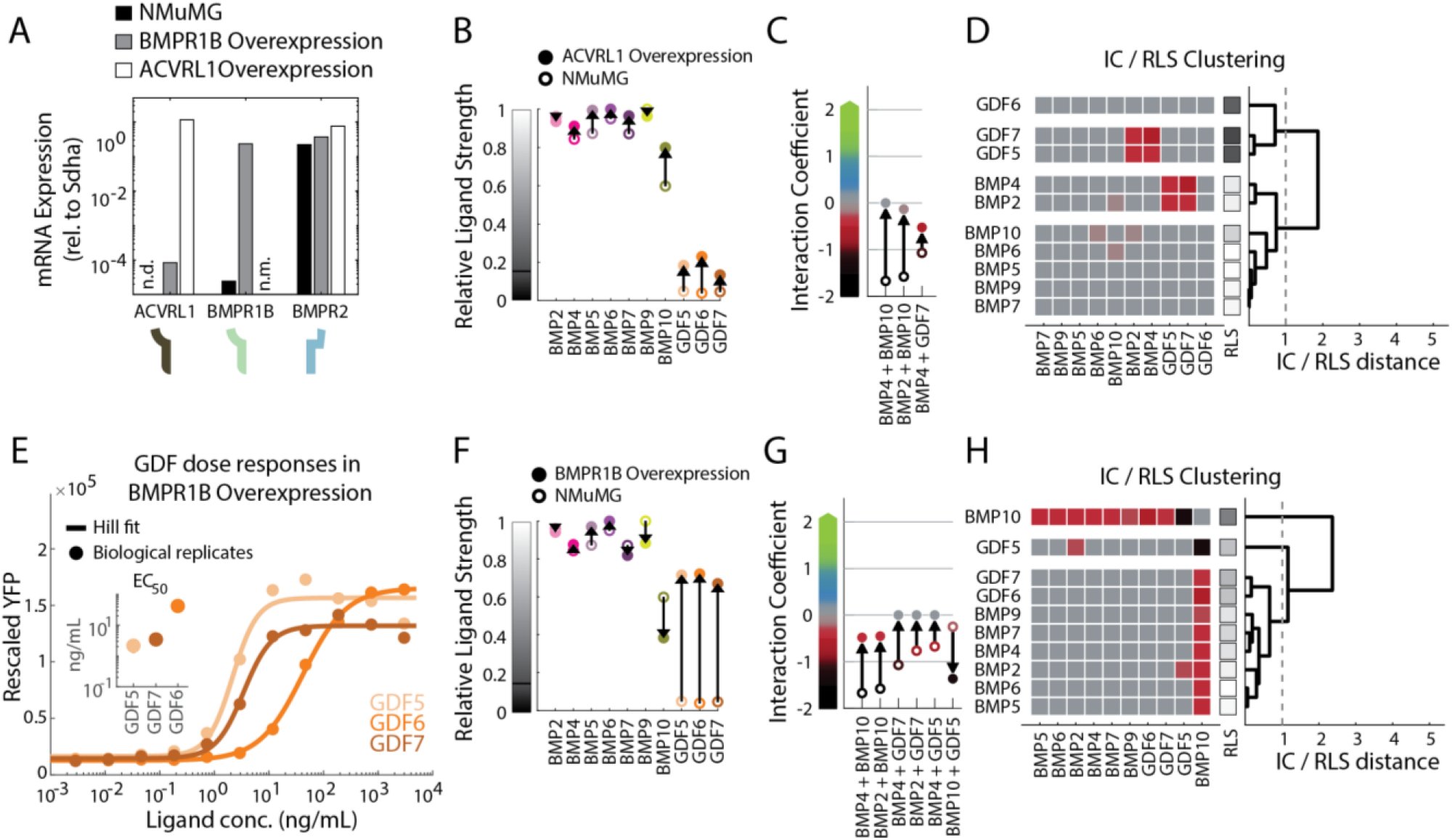
Ectopic expression of missing BMP receptors increases frequency of additive interactions and simplifies ligand equivalence. (A) ACVRL1 and BMPR1B mRNA are not detected (n.d.) or orders of magnitude lower than housekeeping genes in the original NMuMG reporter. Ectopic expression of ACVRL1 and BMPR1B in the NMuMG reporter increases mRNA levels, quantified by RT-qCPR, to match already expressed BMP receptors, such as BMPR2. BMPR1B levels were not measured (n.m.) in cells ectopically expressing ACVRL1. (B) Ectopic ACVRL1 expression (closed circle) in NMuMG cells (open circle) produces small changes in RLS, increasing activation by BMP10, and making GDFs weak activators. (C) Between ectopic ACVRL1 expression (closed circle) and NMuMG cells (open circle), only three ligand pairs show changes in IC larger than 0.5. BMP10’s suppression is lost, while GDF7 more weakly antagonizes BMP4. (D) Hierarchical clustering of all IC and RLS values reveals equivalence groups following ectopic ACVRL1 expression. (E) GDF5, GDF6, and GDF7 activate NMuMG cells following ectopic expression of BMPR1B, though with different EC_50_ values (inset, Hill fit parameters) that are not apparent in other cell lines where GDFs do not activate. (F) Ectopic BMPR1B expression (closed circle) in NMuMG cells (open circle) produces large changes in RLS, converting GDFs into strong activators and slightly decreasing BMP10’s strength. (G) Between ectopic BMPR1B expression (closed circle) and NMuMG cells (open circle), six ligand pairs show changes in IC larger than 0.5, and these pairs involve ligands whose RLS changed significantly. BMP10’s suppression becomes antagonism, but GDF5, rather than antagonizing BMP10, suppresses it. Otherwise, antagonism by GDFs becomes mostly additive. (H) Hierarchical clustering of all IC and RLS values reveals equivalence groups following ectopic ACVRL1 expression. See also Figure 6.

**Figure S7:**
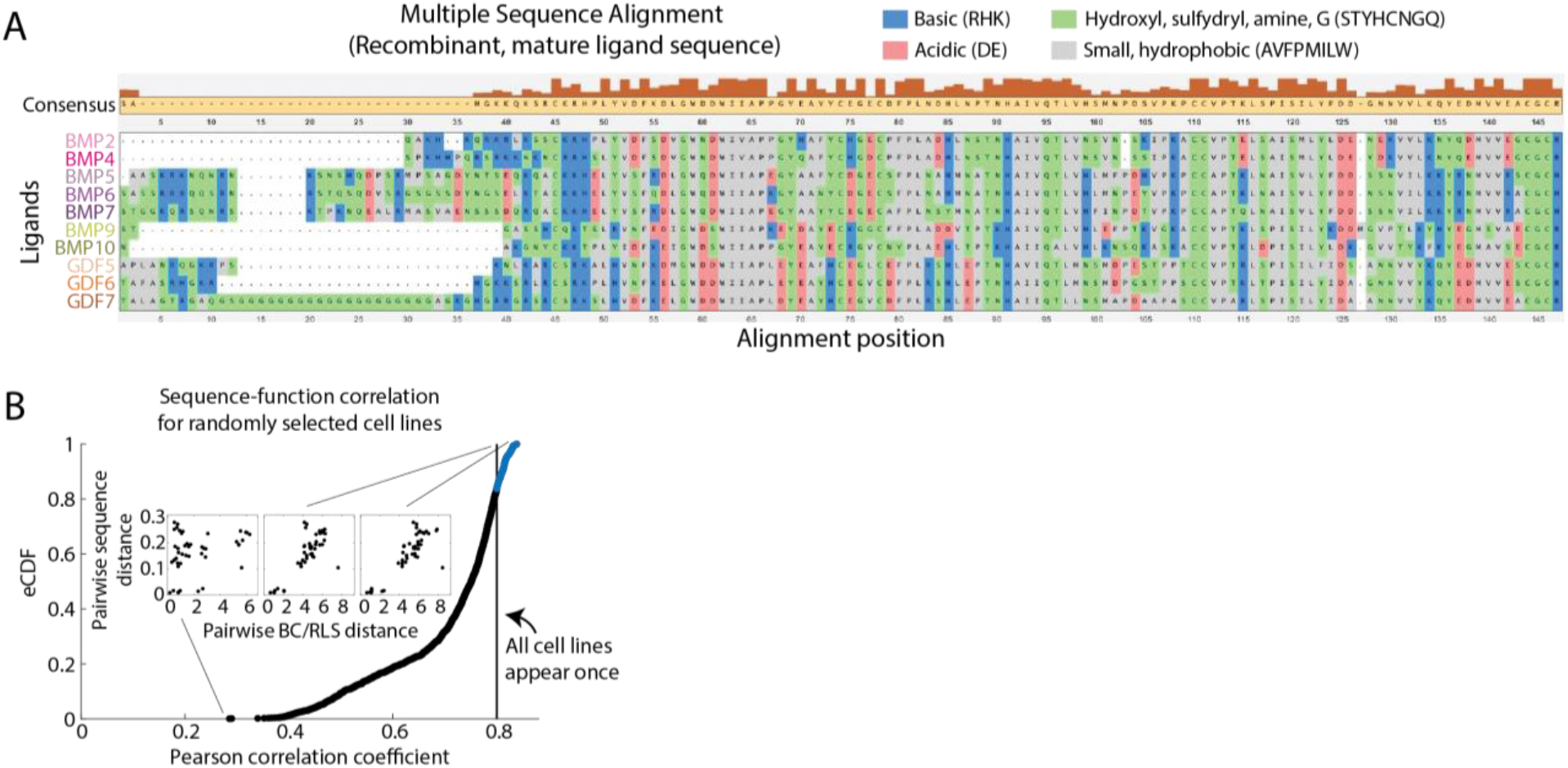
BMP ligand sequence differences correlate with global functional differences. (A) Alignment of recombinant BMP sequences reveals amino acid variations between BMP ligands. (B) The correlation of sequence differences with functional differences was computed for 10,000 random combinations of the seven cell lines. Less than 25% of samples produce higher correlation coefficients than merely including each cell line once. The best and worst correlations, as well as the correlation for each cell line appearing once, are shown in the inset. See also Figure 6.

**Figure S8:**
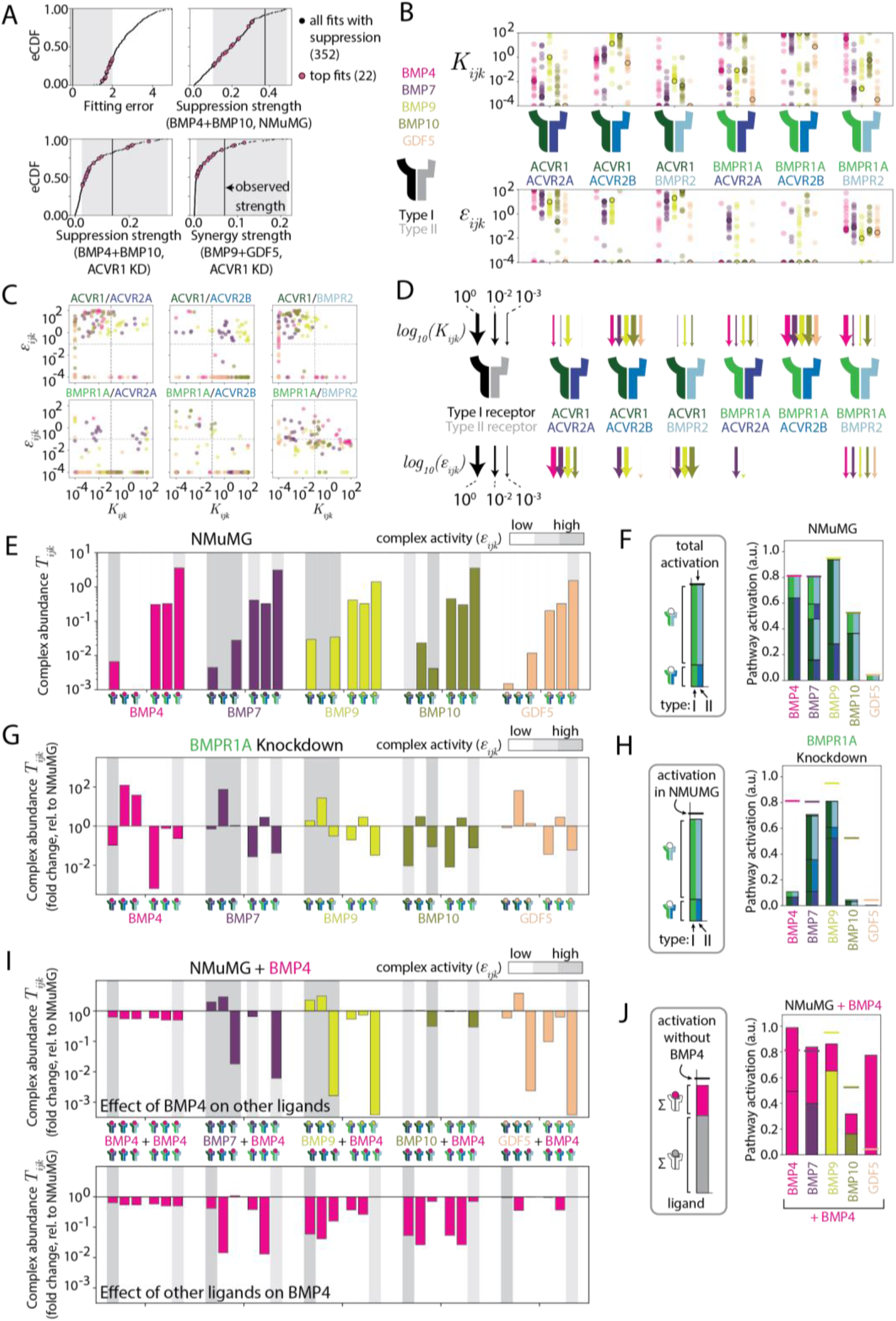
In the model, abundance of high- and low-activity complexes depend on affinities, receptor context, and ligand conext. (A) Of the 6,920 least-squares fits generated, only 352 had fitting error less than 5 while also maintaining the two suppression and synergy interactions observed in the NMuMG and ACVR1 knockdown cells. From these, we selected a top 22 fits which had low error and sufficiently high strength of suppression and synergy. eCDFs for fitting error, suppression strength, and synergy strength are plotted for the 352 solutions which maintained these interactions. The black line shows the observed value, while the gray region shows the accepted regimes for top fits. The pink dots show the top 22 fits which met all four criteria simultaneously. (B) In the top 22 fits, affinity (*K_ijk_*, upper plot) and activity (*ε_ijk_*, lower plot) parameters varied significantly over the allowed range in many cases, such as BMP4’s affinity for BMPR1A-BMPR2. Others were tightly clustered across all parameter sets, such as BMP9’s affinity for ACVR1-BMPR2. The parameter fit shown in Figure 7B is highlighted with black circles. (C) Affinity and activity are anti-correlated across all top fits for certain receptor dimers and ligands, such as ACVR1-BMPR2. (D) A sample parameter set that fits all responses to five ligands in four cell lines are shown as lines for each receptor dimer, where color indicates the ligand species and thickness corresponds to the logarithm of the parameter value. Ligands bind most receptor dimers but activate few. In many cases, affinity and activity are inversely related, as indicated by thick arrows above and thin arrows below, or vice versa. (E) Abundance of each signaling complex (*T_ijk_*) in the mathematical model varies slightly between individual ligands when activating NMuMG cells. The possible complexes are grouped into sets of six, colored by the contained ligand and with the six bars corresponding to the six possible receptor dimers, in the same order as in B and cartooned with the same colors on the x-axis. Grayscale background for each bar indicates the binned (high > 10^0^, low < 10^−2^, medium in between; arbitrary *ε_ijk_* units) activity of that complex. Ligands form similar amounts of the various complexes, but differ in their ability to activate them. (F) Strongly activating ligands achieve similar levels of pathway activation by activating distinct, ligand-specific receptor dimers, whereas weaker ligands activate fewer receptor dimers. Stacked bars show the contribution of various signaling complexes. Left and right halves of each unique bar in the stack are colored to indicate unique pairings of the Type I and Type II receptors, respectively. (G) Bars indicate the fold change, relative to NMuMG cells, in signaling complex abundance following BMPR1A knockdown. Ordering of bars and grayscale background are as in E. BMPR1A knockdown indirectly increases abundance of ACVR2B-containing complexes, which only BMP7 and BMP9 activate strongly. (H) As in D, individual ligands activating the BMPR1A knockdown cell line at saturating concentrations utilize different receptor complexes. Unexpectedly, BMP4 and BMP10 produce lower activity overall, due to the loss of ACVR1-containing complexes that produced most of their pathway activation, while BMP7 and BMP9 shift to other ACVR1-containing complexes. (I) The presence of saturating BMP4 introduces additional, BMP4-containing complexes and changes complex abundance as receptors are shared between the two ligands. Bars indicate the theoretical fold change, relative to ligands signaling alone, in the amount of each receptor complex. For each ligand pair, the full set of complexes are split between the upper plot (the first ligand of the pair) and lower plot (BMP4, the second ligand of the pair). Ordering of bars and grayscale background are as in E. BMP7 and BMP9 lose some activating complexes to BMP4 while also forming new ones, due to their ability to promiscuously activate all ACVR1-containing complexes. (J) Bar height indicates the theoretical contribution of each ligand to overall activation by the ligand pair. Consistent with the effects of competition, overall pathway activation is reduced when BMP4 is combined with BMP10, increased when combined with GDF5, but stays the same, due to sharing of active complexes, when combined with BMP7 and BMP9. See also Figure 7.

